# DNA language models are powerful predictors of genome-wide variant effects

**DOI:** 10.1101/2022.08.22.504706

**Authors:** Gonzalo Benegas, Sanjit Singh Batra, Yun S. Song

## Abstract

The expanding catalog of genome-wide association studies (GWAS) provides biological insights across a variety of species, but identifying the causal variants behind these associations remains a significant challenge. Experimental validation is both labor-intensive and costly, highlighting the need for accurate, scalable computational methods to predict the effects of genetic variants across the entire genome. Inspired by recent progress in natural language processing, unsupervised pre-training on large protein sequence databases has proven successful in extracting complex information related to proteins. These models showcase their ability to learn variant effects in coding regions using an unsupervised approach. Expanding on this idea, we here introduce the **G**enomic **P**re-trained **N**etwork (**GPN**), a model designed to learn genome-wide variant effects through unsupervised pre-training on genomic DNA sequences. Our model also successfully learns gene structure and DNA motifs without any supervision. To demonstrate its utility, we train GPN on *unaligned* reference genomes of *Arabidopsis thaliana* and seven related species within the Brassicales order, and evaluate its ability to predict the functional impact of genetic variants in *Arabidopsis thaliana* by utilizing allele frequencies from the 1001 Genomes Project and a comprehensive database of GWAS. Notably, GPN outperforms predictors based on popular conservation scores such as phyloP and phastCons. Our predictions for *Arabidopsis thaliana* can be visualized as sequence logos in the UCSC Genome Browser (https://genome.ucsc.edu/s/gbenegas/gpn-arabidopsis). We provide code (https://github.com/songlab-cal/gpn) to train GPN for any given species using its DNA sequence alone, enabling unsupervised prediction of variant effects across the entire genome.

---

The emergence of genome-wide association studies (GWAS) has significantly enhanced our ability to examine the genetic basis of complex traits and diseases in both humans and plants. In humans, GWAS have played a crucial role in identifying genetic variants associated with a range of traits, including schizophrenia and obesity [1]. Similarly, in plants, GWAS have shed light on the genetic factors influencing traits such as drought tolerance, disease resistance, and yield [2]. A central challenge in GWAS is pinpointing causal variants for a trait, as linkage disequilibrium (LD) can lead to spurious associations [3]. This process, known as fine-mapping, serves as a foundation for constructing accurate, portable polygenic risk scores and understanding the underlying biological mechanisms. Although experimental validation of causal variants is the gold standard, it is not scalable. Instead, a scalable fine-mapping strategy involves utilizing computational variant effect predictors [4], which vary from conservation scores to deep learning models trained on functional genomics data. Accurate variant effect prediction is also vital for diagnosing rare diseases and interpreting rare variants that lie beyond the scope of traditional GWAS [5].

Recently, state-of-the-art performance in predicting the effects of missense (coding) variants has been achieved by training unsupervised models on extensive protein sequence databases [6] or their corresponding multiple sequence alignments [7]. These large language models can predict missense variant effects in an unsupervised manner, without the need for additional training on labeled data. This progress has been driven by advancements in natural language processing, where significant strides have been made by pre-training language models on vast text corpora. Pre-trained models such as BERT can be fine-tuned for downstream tasks such as sentiment analysis [8]. More recently, language models like GPT-4 have demonstrated impressive leaps in test performance across various disciplines, from law to computer science [9].

A widely-used approach to interpreting non-coding variant effects involves training a supervised model to predict functional genomics data — such as chromatin accessibility, transcription factor binding, or gene expression — and then evaluating variants based on how they disrupt these predictions. This approach was first introduced by DeepSEA [10], which utilized 919 functional genomics tracks, and has since been refined by Enformer [11] with 6,956 tracks and Sei [12] with 21,907 tracks. However, this approach’s success depends on the availability of high-quality functional genomics data from a diverse array of cell types, which can be prohibitively expensive to generate for most species. Certain models focus on specific classes of non-coding variants. For instance, classifiers trained solely on sequence data can predict the impact of intron variants on splicing patterns [13, 14]. To evaluate the effects of regulatory variants, Lee *et al*. [15] developed a support vector machine that distinguishes putative regulatory sequences from random genomic sequences. More recently, a deep learning model capable of predicting Hi-C signal from sequence data demonstrated its potential to predict the impact of regulatory variants on DNA folding within the nucleus [16]. Additionally, a deep learning model [17] was successfully trained to predict DNA methylation levels of CpG sites from sequence data, enabling the prediction of non-coding variant effects on DNA methylation.

However, variant type-specific models are inherently not well-suited for diagnosing rare diseases, fine-mapping, or calculating polygenic risk scores, as these tasks necessitate the comparison of genome-wide variants all together. For instance, a model that is exclusively designed for either missense or regulatory variants would not be able to prioritize between a *de novo* missense variant and a *de novo* promoter variant observed in a patient with a rare disease. An important class of genome-wide scores are conservation scores such as phyloP [18] and phastCons [19], which are computed from genome-wide alignment of multiple species. Since these do not require functional genomics data, they have been widely applied to many systems, including non-model organisms [20]. In humans, CADD is another important genome-wide variant effect predictor that combines conservation and functional genomics annotations, and is trained to distinguish between an inferred set of putative benign and putative pathogenic variants [21, 22].

In this paper, we introduce the **G**enomic **P**re-trained **N**etwork (**GPN**), a multi-species DNA language model trained using self-supervision. While existing DNA language models [23, 24, 25, 26, 27, 28, 29] have not yet demonstrated the ability to make accurate variant effect predictions based on self-supervision alone, GPN presents a unified approach capable of accurate unsupervised prediction of genome-wide variant effects. We demonstrate its utility by achieving state-of-the-art performance in *Arabidopsis thaliana*, a model organism for plant biology closely related to many agriculturally important species, as well as a source of insight into human diseases [30]. Moreover, GPN outperforms genome-wide conservation scores such as phyloP and PhastCons, which rely on whole-genome alignments of 18 closely related species [20]. GPN’s internal representation of DNA sequences can distinguish genomic regions like introns, untranslated regions, and coding sequences. Additionally, the confidence of GPN’s predictions can help reveal regulatory grammar, such as transcription factor binding motifs. Our results lay the foundation for developing state-of-the-art genome-wide variant effect predictors for any species using genomic sequence alone, which can be readily integrated into GWAS fine-mapping and polygenic risk scores.

## Results

### Training a multi-species DNA language model

We used *unaligned* reference genomes from *Arabidopsis thaliana* and seven related species within the Brassicales order to pre-train a language model based on a convolutional neural network (Supplementary Table S1). This model was designed to predict masked nucleotides conditioned on their local genomic context (Figure 1, *Materials and Methods*). During the training process, we encountered challenges with repetitive elements, which can be functionally significant but are heavily overrepresented in the genomes [31]. We found that reducing the weight of prediction loss for repetitive regions led to improved test perplexity in non-repetitive regions, which are often of greater interest (Supplementary Table S2). Compared to full down-weighting, moderate down-weighting results in a similar improvement in perplexity for non-repetitive regions without sacrificing genome-wide perplexity as much. Consequently, we focus on this model throughout the remainder of the paper unless otherwise specified.

**Figure 1:**
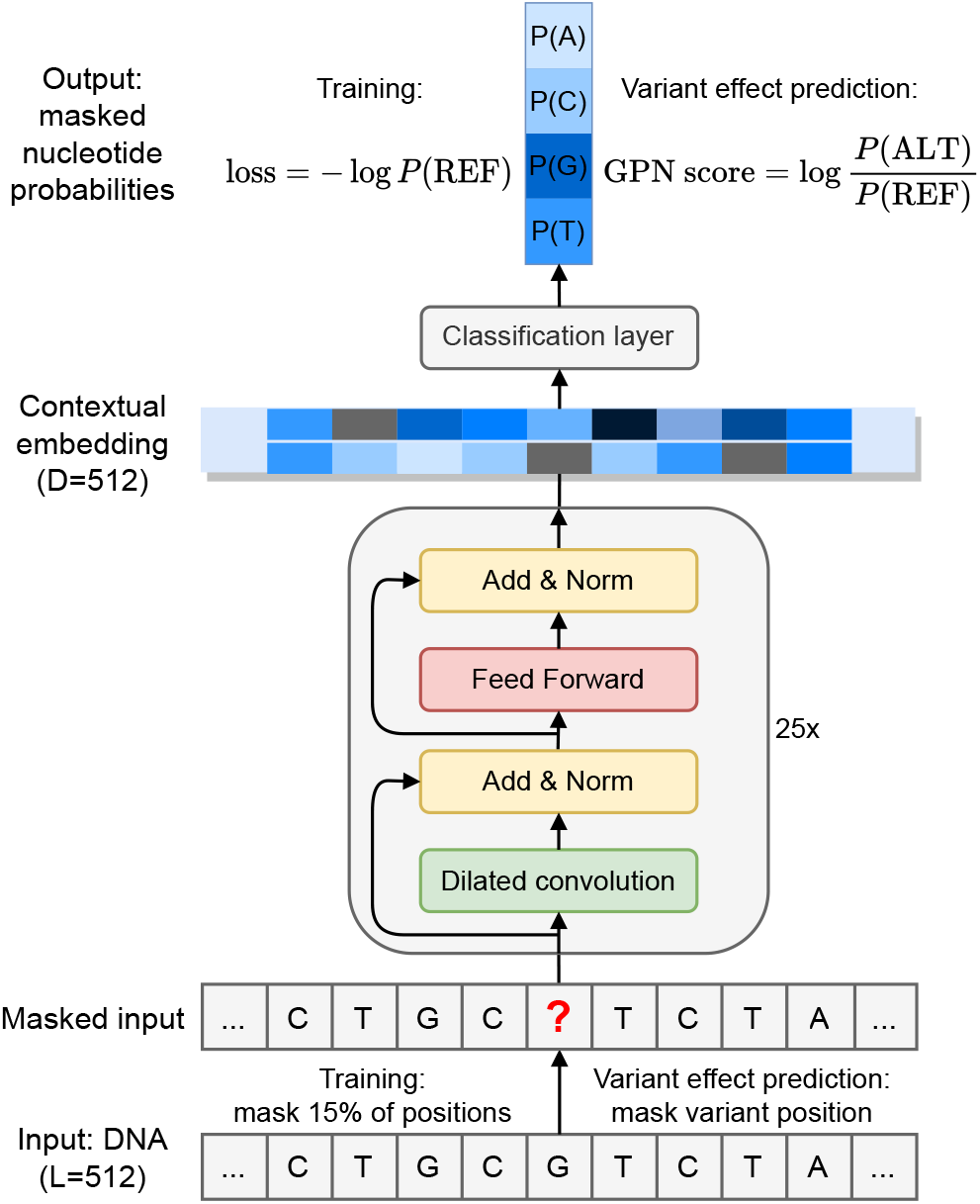
Overview of GPN (Genomic Pre-trained Network). The input is a 512-bp DNA sequence where certain positions have been masked, and the goal is to predict the nucleotides at the masked positions. During training, 15% of the positions are masked. During variant effect prediction, only the variant position is masked. The sequence is processed through a convolutional neural network resulting in a high-dimensional contextual embedding of each position. Then, a final layer outputs four nucleotide probabilities at each masked position. The model is trained on the reference sequence with the cross-entropy loss. The GPN variant effect prediction score is defined as the log-likelihood ratio between the alternate and reference allele. L: window length in base pairs. D: embedding dimension. REF: reference allele. ALT: alternate allele.

### Unsupervised clustering of genomic regions

To understand how well the model has learned the structure of the genome, we averaged GPN’s contextual embeddings (512 dimensions) of nucleotides over 100 base pair (bp) windows from the reference genome and visualized them using UMAP [32] (Figure 2a). Notably, GPN, trained without any supervision, has learned to distinguish genomic regions such as intergenic, introns, coding sequences (CDS), untranslated regions (UTR) and non-coding RNA (ncRNA). To quantify GPN’s ability to distinguish genomic regions, we trained a logistic regression classifier using the averaged embeddings as features, achieving the highest accuracy on CDS (96%) and the lowest on ncRNA (51%), the least frequent class. As summarized in Figure 2b, the highest confusion was observed between intergenic regions and ncRNAs; this may be partly explained by errors in ncRNA annotation, which is especially challenging given their low expression levels and poor conservation [33]. This level of classification accuracy cannot be achieved merely through *k*-mer frequencies (Supplementary Figure S1). We also note that, to some extent, GPN embeddings can distinguish different repeat families (Supplementary Figure S2).

**Figure 2:**
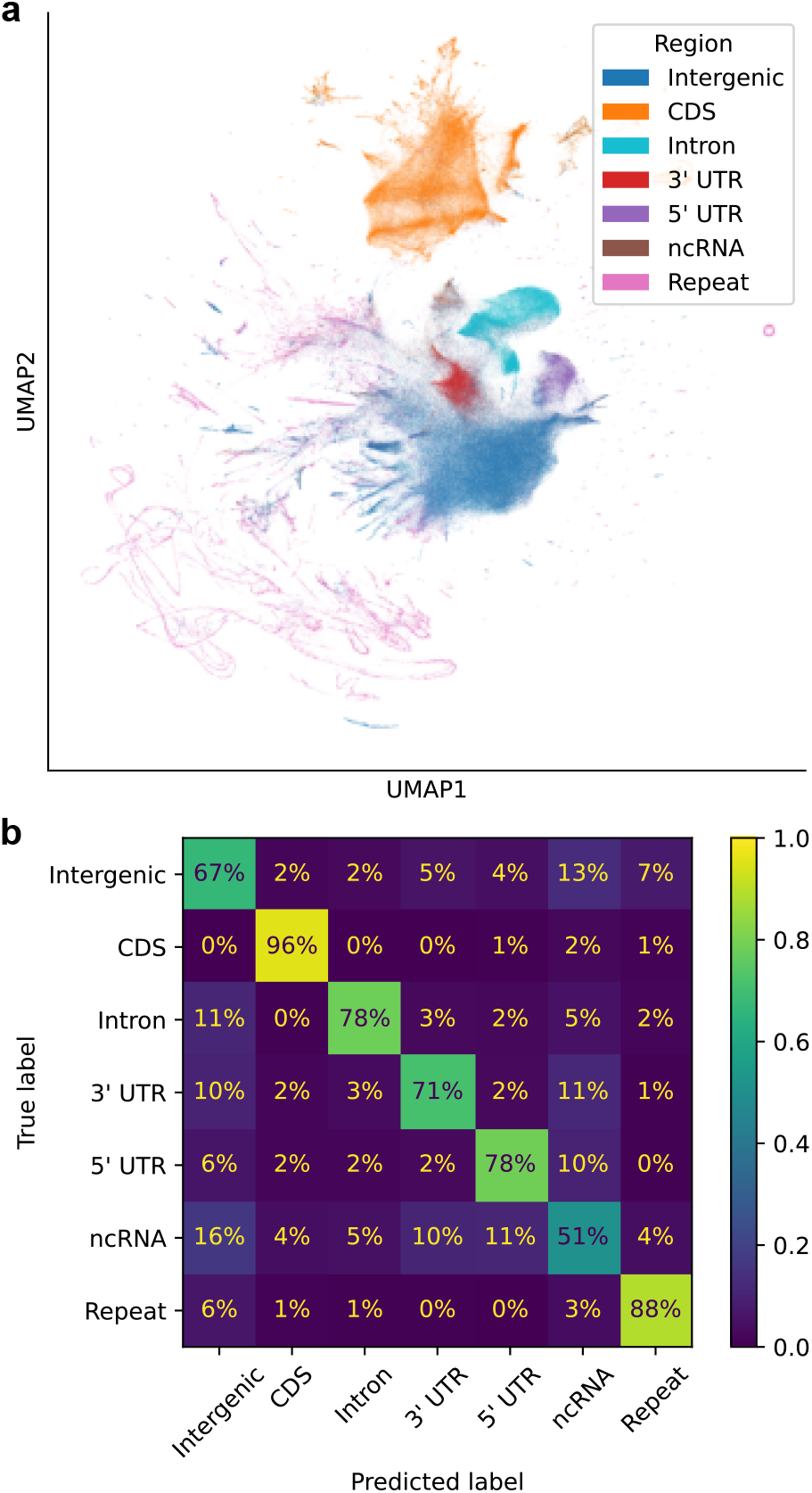
Unsupervised clustering of genomic windows. (a) UMAP visualization of GPN embeddings averaged over non-overlapping 100 bp windows along the genome, annotated with gene region. (b) Confusion matrix for classification of gene regions using a logistic regression model trained on averaged embeddings. Each chromosome was predicted from a model trained on the remaining chromosomes.

### DNA motifs revealed by high-confidence model predictions

To further understand GPN, we individually masked each position in the genome and obtained the model output distribution over nucleotides, given its context. To facilitate utilizing these predicted distributions, we created sequence logos that can be visualized in the UCSC Genome Browser [34, 35] (https://genome.ucsc.edu/s/gbenegas/gpn-arabidopsis), where the height of each letter is proportional to its probability, and the overall height is given by the information content, measured in bits [36] (see Figure 3a for an example). The model’s prediction confidence correlates with the expected functionality of the sites. For example, exonic positions are predicted with higher confidence than the surrounding introns, except for the canonical splice acceptor and donor dinucleotide motifs. Similarly, the third nucleotide position within codons, which usually does not affect amino acid identity, is generally predicted with lower confidence than the first two positions. Start and stop codon motifs are also generally well predicted (examples in Supplementary Figure S3). Across the test chromosome, model perplexities in splice acceptors (median = 1.03), splice donors (median = 1.02), start codons (median = 1.10), and stop codons (median = 2.81) are significantly smaller than those in intergenic and intronic regions (median = 3.14, all Mann–Whitney *p*-values *<* 10^*−*239^, Supplementary Figure S4).

**Figure 3:**
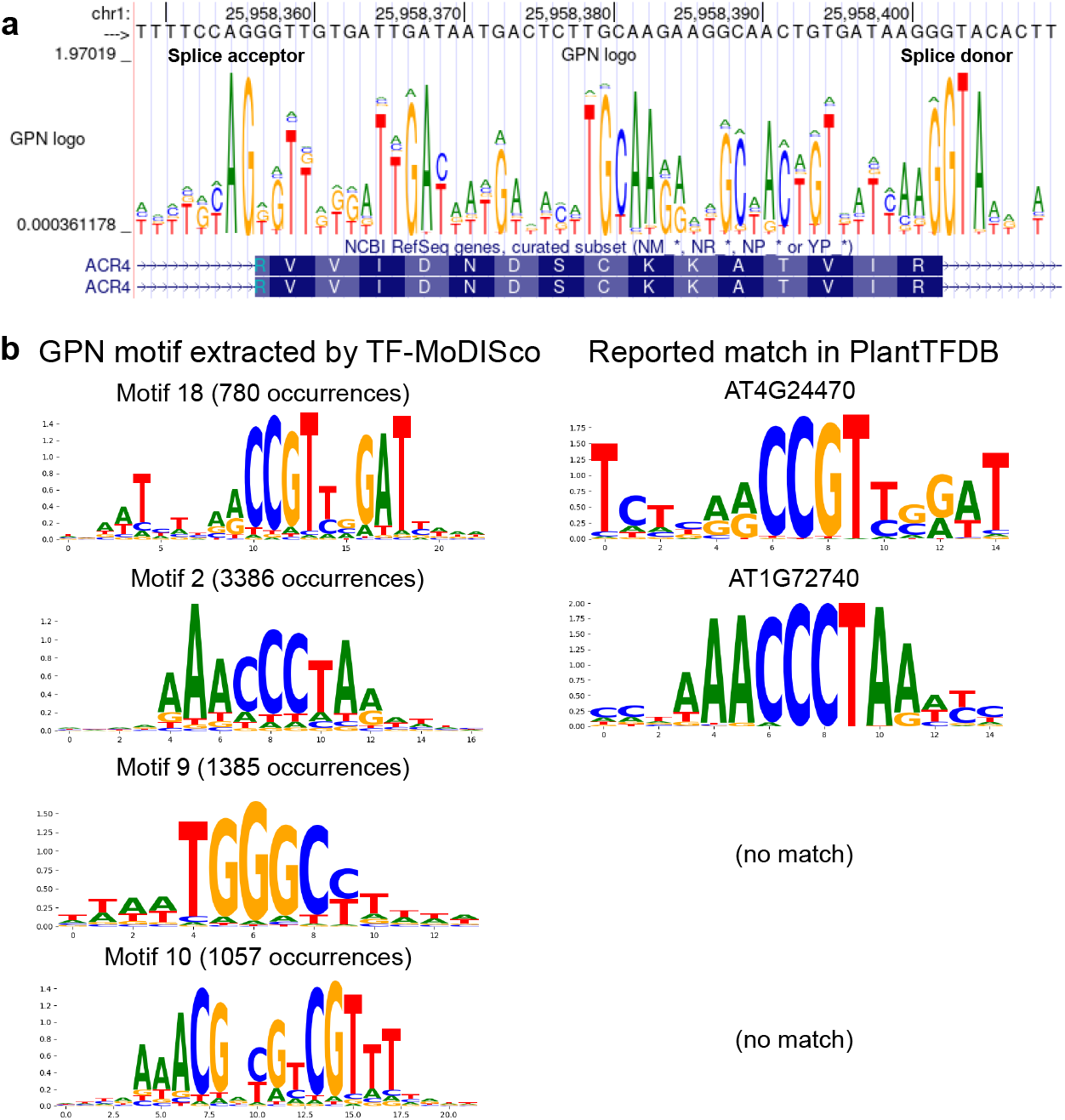
Sequence logos derived from model predictions. Each position in the genome was independently masked and the model distribution over the four nucleotides was computed. (a) Sequence logo visualized in the UCSC Genome Browser (https://genome.ucsc.edu/s/gbenegas/gpn-arabidopsis). The height of each letter is proportional to its probability, while the overall height at each position is equal to 2 minus the entropy of the distribution. (b) Example GPN motifs in promoter regions, extracted by TF-MoDISco, with significant matches in PlantTFDB.

We hypothesized that scanning promoters for small regions of high-confidence GPN predictions could help identify transcription factor binding sites. To achieve this, we adapted TF-MoDISco [37], a tool for *de novo* discovery of transcription factor binding sites using supervised models. This tool clusters high-scoring regions into motifs and compares them to databases of known motifs. Applying the adapted TF-MoDISco to GPN scores in promoter regions, we discovered approximately a hundred and sixty motifs (Supplementary Figure S5), with four examples shown in Figure 3b, the first two having a significant match in PlantTFDB [20]. Some of the discovered motifs are well-documented in the literature but do not have a significant match in this database, such as the third motif [38] in Figure 3b. Some motifs could represent novel promoter elements, like the fourth motif, which is palindromic with symmetrical entropies, suggesting that it could potentially form RNA or DNA alternative secondary structure [39].

### Unsupervised variant effect prediction

GPN can be employed to calculate a pathogenicity or functionality score for any single-nucleotide polymorphism (SNP) in the genome using the log-likelihood ratio between the alternate and reference allele (GPN score, Figure 1). Visually, this involves comparing the heights of the letters in the logo plot (Figure 3a).

#### In silico mutagenesis

We first computed GPN scores for *in silico* mutagenesis of SNPs within a 1 Mb region and aggregated the results across variant types (Figure 4). The ranking of variant types based on the lowest percentile of GPN scores is generally consistent with established notions of deleteriousness [40]^1^. For example, the four lowest scored variant types are splice donor, splice acceptor, stop gained and start lost variants, which significantly disrupt the open reading frame. As expected, missense variants are ranked before synonymous variants. However, we observed that some variants within repetitive elements were assigned rather low GPN scores, ranking close to missense variants. Furthermore, the proportion of low GPN scores for repeat variants depends on the training loss weight on repeats (Supplementary Figure S5a). More precisely, in models with 0.0 and 0.1 down-weighting, respectively, 8% and 9% of repeat variants are ranked before the first decile of missense variants. These represent a substantial decrease compared to the 27% observed in the model without any down-weighting (Supplementary Figure S5b, Fisher’s exact test *p <* 10^*−*300^).

**Figure 4:**
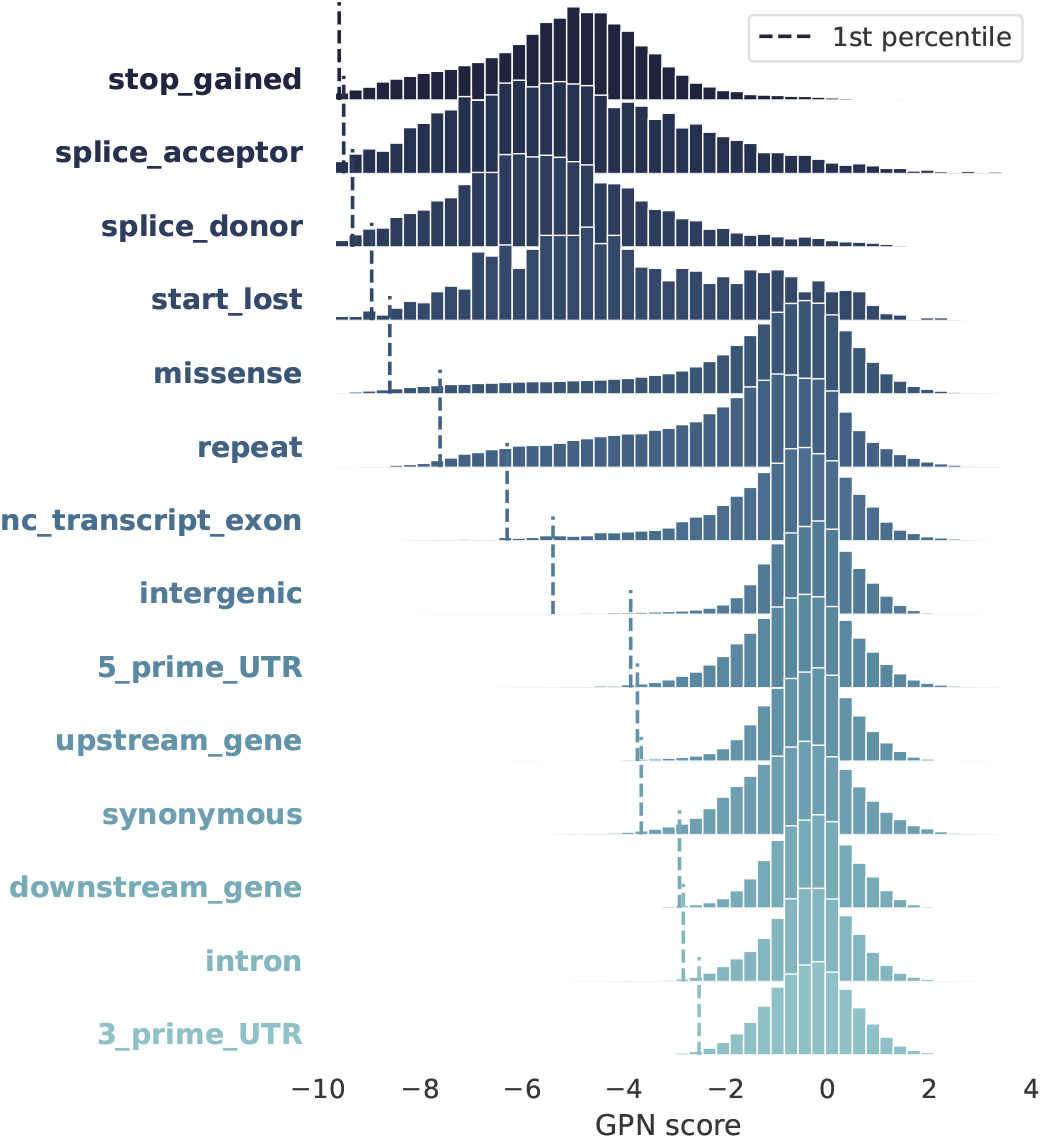
Variant effect prediction: *in silico* mutagenesis. Distribution of GPN scores computed for all possible single-nucleotide polymorphisms (SNPs) in a 1 Mb region, across categories, sorted by 1st percentile (dashed vertical lines).

#### Benchmarking using allele frequencies in 1001 Genomes

Following our *in silico* mutagenesis experiments, we analyzed over 10 million SNPs from naturally occurring accessions of the 1001 Genomes Project [41]. While most variants have a neutral GPN score, there is a heavy tail of putative functional variants with negative GPN scores (Figure 5a). Notably, variants with lower GPN scores are, on average, less frequent in the population, suggesting they could be under purifying selection (Figure 5b, full distribution in Supplementary Figure S6). To evaluate the capability of identifying putative functional variants, we assessed the enrichment of rare versus common variants in the tail of genome-wide score distributions. Putative functional SNPs, defined as the lowest 0.1% of GPN scores, exhibit a 5.5-fold enrichment in rare variants (Figure 5c); see Supplementary Figure S7 for different allele frequency thresholds. GPN outperforms other genome-wide variant effect predictors for *Arabidopsis*, specifically phyloP and phastCons, which are conservation scores derived from a broader set of 18 Brassicales species (Figure 5d). In fact, GPN scores are only weakly correlated with phyloP (*r* = 0.22, *p <* 10^*−*300^) and phastCons (*r* = 0.13, *p <* 10^*−*300^). We also considered the alternative abs(phyloP) (the absolute value of phyloP), but it did not achieve a significant enrichment. A notable advantage of GPN is that it is able to score variants that could not be scored by phyloP and phastCons due to unsuccessful whole-genome alignment (14.2% of all variants). GPN performs comparably to phyloP and phastCons when using less stringent thresholds for defining putative functional SNPs (Supplementary Figure S8), indicating its particular strength in detecting deleterious variants at the extreme tail. GPN also achieves significant odds ratios when computed only within particular variant classes, but its performance relative to phyloP and phastCons varies (Supplementary Figure S8). On a separate note, a slightly higher odds ratio is achieved by the GPN model trained with an intermediate loss weight on repeats (Supplementary Figure S5c). The model trained on only a single species performs substantially worse (Supplementary Figure S9a).

**Figure 5:**
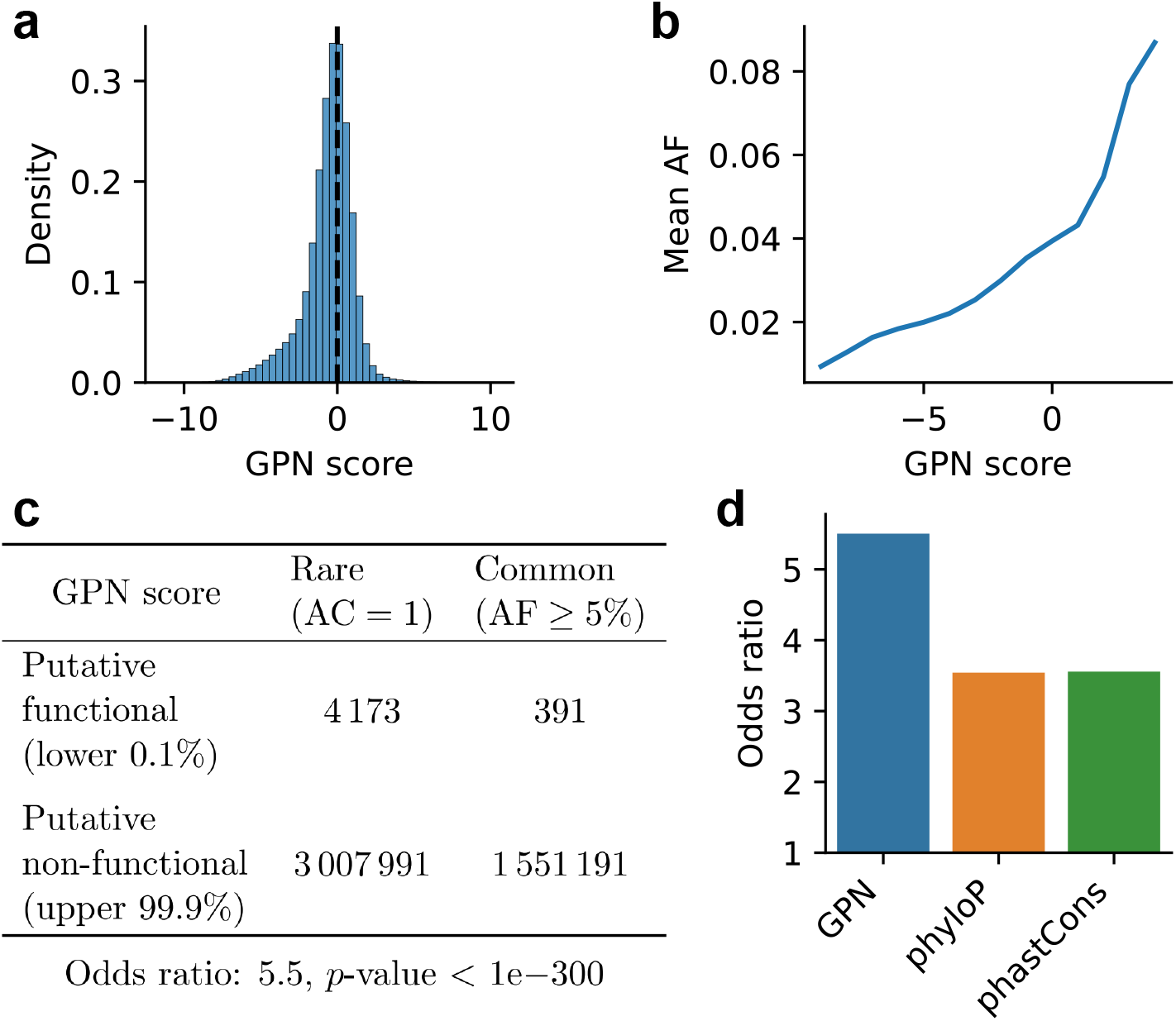
Variant effect prediction: rare vs. common. The GPN score was computed for over 10 million variants in the 1001 Genomes. (a) Distribution of GPN scores. (b) Mean allele frequency for different GPN score bins ([*−*9.5, *−*8.5), [*−*8.5, *−*7.5), …, [3.5, 4.5)). (c) Contingency table and odds ratio showing enrichment of putative functional GPN scores in rare variants. AC: allele count. AF: allele frequency. (d) Comparison of odds ratios as in (c) obtained with different models. abs(phyloP) is excluded as it did not achieve a significant enrichment.

#### Enrichment of GWAS hits in regions with low GPN scores

In our pursuit to further evaluate the efficacy of GPN, we examined the AraGWAS Catalog [42], a comprehensive database of genomewide association studies (GWAS) in *Arabidopsis thaliana*. We hypothesized that GWAS hits may be enriched in regions with low GPN scores. An advantage of GPN is that it can give substantially different scores to variants in strong linkage disequilibrium (LD) with each other, if their surrounding contexts are different (e.g., see Figure 6a, top). In contrast, the standard GWAS would give similar scores to such variants; in particular, neutral variants in strong LD with a functional variant would also be associated with a trait. To account for this difference, we devised a new score, GPN*×*LD, which weighs GPN scores by LD (*Materials and Methods*). With this approach, GPN*×*LD effectively distinguishes GWAS hits from non-hits in this example locus (Figure 6a, bottom). More generally across the genome and all traits, the tail of GPN*×*LD scores is greatly enriched in GWAS hits, much more so than the tail of raw GPN scores (Figure 6b). In particular, by analyzing odds ratios (Figure 6c), we found that SNPs with the lower 1% of GPN*×*LD scores are 10.3-fold enriched in GWAS hits compared to the upper 99% of GPN*×*LD scores, while less than 7.5-fold enrichment was observed for other methods (Figure 6d); see Supplementary Figure S10 for different thresholds. Using the Bonferroni correction instead of the permutation-based significance threshold recommended by AraGWAS [43] yields lower odds ratios for all methods, but GPN*×*LD still achieves the highest enrichment (Supplementary Figure S11). Interestingly, the GPN model trained with an intermediate loss weight on repeats achieves the best performance (Supplementary Figure S5d). The model trained on only a single species performs worse (Supplementary Figure S9b). Furthermore, GPN*×*LD achieves much higher odds ratios when considering the full variant set, including regions that do not align to other Brassicales (Figure 6e); failed alignment could be partly due to genomic rearrangements that may be potentially associated with local adaptation in *Arabidopsis thaliana* [44].

**Figure 6:**
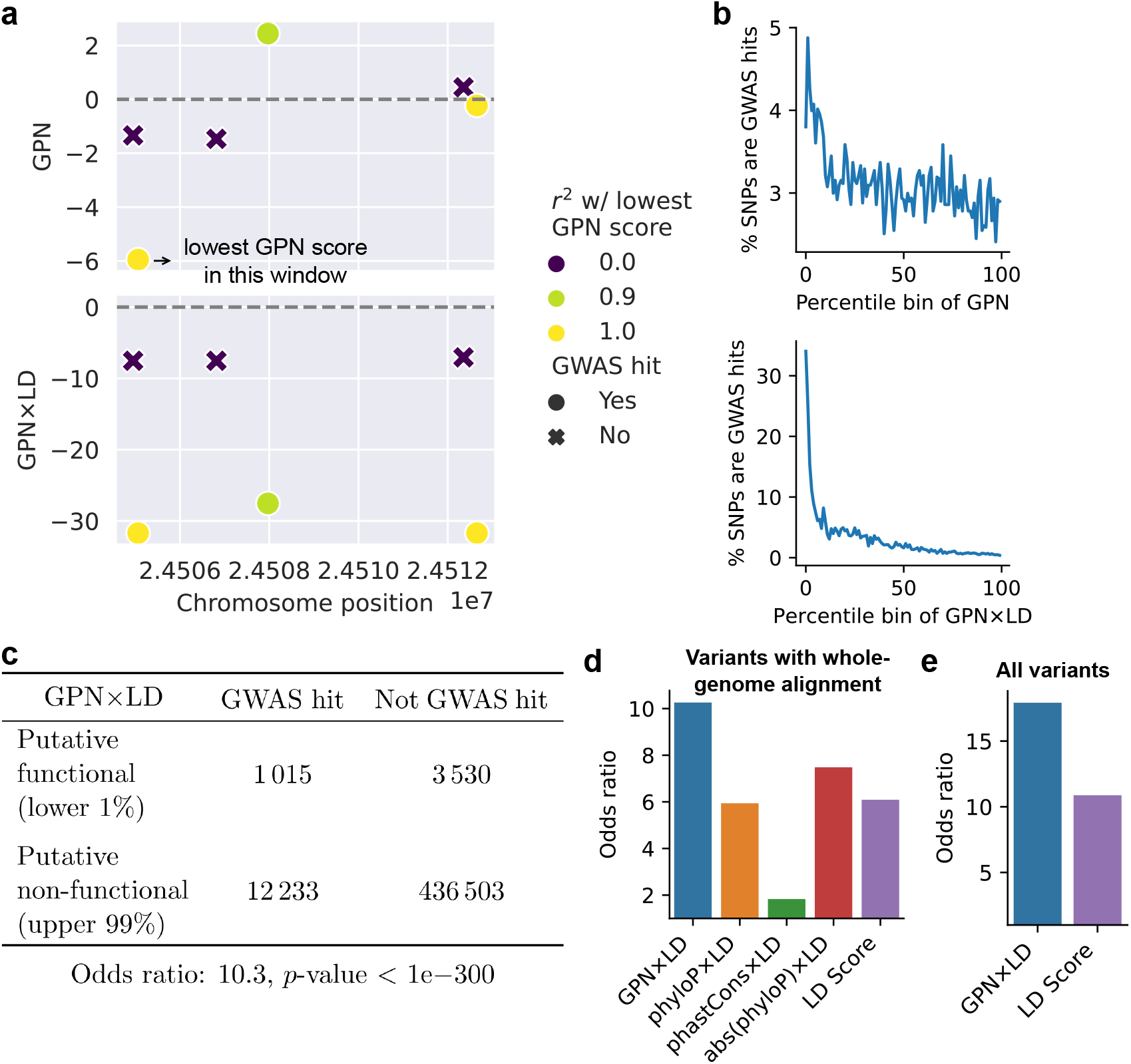
Variant effect prediction: GWAS. GPN scores were analyzed for around half a million variants tested in AraGWAS. (a) Example window with six variants tested for association with maximum temperature in January. GPN*×*LD successfully separates GWAS hits and non-hits. (b) Percentage of GWAS hits (for any trait) in each percentile bin of GPN and GPN*×*LD scores. (c) Contingency table and odds ratio showing enrichment of GWAS hits (for any trait) in putative functional (associated) GPN*×*LD scores. (d) Comparison of odds ratios obtained with different models (*n* = 453, 281 variants with whole-genome alignment). (e) Odds ratios with the full variant set (*n* = 510, 462).

## Discussion

Here we present the first unsupervised genome-wide variant effect predictor based on unsupervised pre-training of DNA language models. We demonstrate that GPN outperforms other genome-wide variant effect predictors in *Arabidopsis thaliana*, a model species for plant biology. Since GPN is trained only on DNA sequence, it can be readily applied to understudied non-model organisms even in the absence of extensive functional genomics data, while still providing state-of-the-art unsupervised variant effect prediction genome-wide.

We can think of GPN as a generalized conservation score. Similar to phyloP and phastCons, GPN is genome-wide, can be trained on genomic sequence alone, and is cell-type and mechanism agnostic [45]. The key distinction is that while phyloP and phastCons only consider nucleotide frequencies at a specific site, GPN can learn from joint nucleotide distributions across all similar contexts appearing in the genome. Furthermore, GPN does not rely on whole-genome alignments, which can often have a lower quality in non-coding regions.

The capability of GPN to score genome-wide variants on a unified scale renders it ideal for integration into rare disease diagnosis, fine-mapping, and polygenic risk scores, including burden tests. The separation of genomic regions based on GPN embeddings suggests that it could be further fine-tuned for *de novo* genome annotation. Combining GPN predictions with TF-MoDISco offers a promising strategy for discovering functional motifs. Although in this study we focused on transcription factor binding sites, we believe that GPN predictions around splice junctions could also facilitate the identification of splicing factor binding sites.

Repetitive elements, which are inherent components of eukaryotic genomes, pose several challenges that have been underexplored in DNA language modeling studies. First, these elements are significantly over-represented [31]. The improved perplexity in non-repetitive regions upon down-weighting repeats can be attributed to the model allocating fewer parameters exclusively to repetitive elements. Second, repetitive elements display reduced sequence variation compared to other regions, in particular younger repeats with little time to accumulate mutations [46]. We believe that these factors together may cause differences in model likelihoods in these regions to be less clearly associated with differences in fitness. Our proposed down-weighting of repeats only partially mitigates these issues, and we encourage further investigation by the scientific community. Potential research directions include examining the effects of down-weighting repeats based on their respective families or inferred age.

While the current implementation of GPN achieves state-of-the-art variant effect prediction for *Arabidopsis thaliana*, there is room for improving its training scheme. Mounting evidence suggests that larger models and more extensive training data can enhance performance [47]. Our current proof-of-concept model is considerably smaller — by 200 times — than the largest published protein language model [48]. One strategy to improve GPN, inspired by protein modeling, involves explicitly incorporating multiple sequence alignments [49, 50]. However, this enhancement will be bottle-necked by the quality of alignment in non-coding genome regions. Other promising avenues for DNA language modeling include incorporating DNA-specific inductive biases, such as reversecomplement equivariance [51], as opposed to our current method of averaging model outputs for both strands during testing. Additionally, integrating long-range information using recent advances in state space models [52] may further boost performance. In conclusion, DNA language models represent a powerful framework for genome-wide variant effect prediction, and we believe that exploring the above avenues to further improve GPN would be worthwhile.

## Materials and Methods

### Pre-training

We obtained a list of Brassicales reference genome assemblies from NCBI Genome (https://www.ncbi.nlm.nih.gov/data-hub/genome/), filtered for RefSeq-annotated and kept only one per genus, resulting in a total of 8 reference genomes (Supplementary Table S1). We held out *Arabidopsis thaliana* chromosomes 4 and 5 for validation and testing, respectively. For each genome, we subsampled genomic windows of size 512 bp, with a step of 256 bp and augmented with the reverse complement. However, we did not draw genomic windows uniformly from the whole genome, but emphasized certain regions. In particular, we took the union of exons (with a small intronic flank), promoters (1000 bp upstream of transcription start sites) as well as an equivalent amount of random windows from the whole genome. We think this decision may improve performance, but leave experimentation for further studies. Additionally, we subset the number of windows from each genome to the number of windows from *Arabidopsis*, given its unusually small genome.

We set up a masked language modeling task [8], in which 15% of the tokens in a nucleotide sequence were masked and had to be predicted from their context. In contrast to most DNA language models that tokenize sequences into overlapping *k*-mers [24, 26, 28] or use byte-pair encoding [23], we used bare nucleotides as tokens. While a thorough benchmark of different tokenization strategies is lacking, using single-nucleotide tokens makes interpretation easier, in particular for unsupervised variant effect prediction.

While language model pre-training successes were first showcased by transformer architectures, convolutional models have shown similarly good performance in natural language [53] and protein modeling [54]. In our initial experiments, we noticed that convolutional models converged faster than transformer models. The locality of convolutions may be a good inductive bias for modeling DNA sequences at this scale. The linear complexity of convolution also simplifies inference or finetuning on longer sequences such as entire chromosomes, which in the case of transformers might require chunking (with some overlap) and aggregating the results.

We implemented GPN, a convolutional neural network, using the Hugging Face library [55]. The masked DNA sequence was one-hot encoded and then consecutively processed by 25 convolutional blocks. Each convolutional block consisted of a dilated convolutional layer followed by a feed-forward layer, with intermediate residual connections and layer normalization (Figure 1). Throughout the layers, the embedding dimension (number of convolutional filters) was kept fixed at 512. The dilation was increased exponentially up to a certain value and then cycled. A list of hyperparameters is displayed in Supplementary Table S3. We trained three models varying only in the loss weight on repetitive elements (marked lowercase in the fasta file). We trained each model for 150 K steps, taking approximately 4 days with 4 NVIDIA A100 80GB GPUs. Perplexity is defined as the exponentiation of the cross-entropy loss, which is equivalent to 1 over the probability given to the correct nucleotide. Test perplexity is displayed in Supplementary Table S2. We also trained a separate model on the single genome of *Arabidopsis thaliana*, with a repeat weight of 0.1 and the same hyperparameters except for only 12,000 steps with decaying learning rate, as we noticed it would soon start overfitting. This model obtained a higher test perplexity of 3.13 (3.17 on non-repeat regions).

### Analysis of model embeddings

Model embeddings were averaged over non-overlapping 100-bp windows. Embeddings from the forward and reverse strand were averaged, and then standardized. UMAP was run with default parameters. The gene annotation was downloaded from EnsemblPlants. The annotation of repetitive elements was downloaded from http://ucsc.gao-lab.org/cgi-bin/hgTables?hgsid=167291_E9nY5UIAQRUOAR01xJAsum4vDukw. We considered intergenic regions with 100% overlap with repeats as a separate “Repeat” class. Windows with ambiguous annotation (e.g., 50% CDS and 50% intron) were excluded from the analysis. Genomic region classification was performed with logistic regression as implemented by scikit-learn [56], using class weight inversely proportional to frequency and L2 regularization strength chosen via cross-validation. Windows in each chromosome were predicted by a model trained on the remaining chromosomes.

### Motif analysis

Each position in the genome was independently masked and the model distribution over nucleotides was extracted. The distribution was averaged between the results from the forward and reverse strands. The held-out model perplexity was computed for splice acceptors, splice donors, start codons, stop codons, and 216 224 positions randomly sampled from intergenic and intronic regions, after excluding repeats.

An adaptation of TF-MoDISco was run with model predictions in regions 1000 bp upstream and downstream of transcription start sites, after filtering repeats and coding exons. The exact score fed into Modisco was the nucleotide probability minus 0.25, so it would be roughly centered at 0. Since TF-MoDISco expects genomic windows of equal length, we concatenated our variable-length windows into one large window, interspersed with 20 undefined ‘N’ nucleotides.

### Variant effect prediction

We scored variants by masking the position and calculating the log-likelihood ratio between the alternate and reference allele. Scores computed from the forward and reverse strands were averaged. We calculated odds ratio and *p*-value with Fisher’s exact test. When comparing to phyloP and phastCons, we excluded variants where these scores are undefined (due to the lack of whole-genome alignment).

All possible SNPs in the region Chr5:3,500,000-4,500,000 were generated and their consequences annotated with Ensembl Variant Effect Predictor [40] web interface https://plants.ensembl.org/Arabidopsis_thaliana/Tools/VEP, with the upstream/downstream argument set to 500, used to call variants as upstream/downstream instead of intergenic. We compared scores for variant types with at least 1000 variants, and we excluded variants with different consequences in different transcripts.

The 1001 Genomes genotype matrix was downloaded from https://aragwas.1001genomes.org/api/genotypes/download and combined with metadata from https://1001genomes.org/data/GMI-MPI/releases/v3.1/1001genomes_snp-short-indel_only_ACGTN.vcf.gz. This genotype matrix is binary, since all the accessions are homozygous, as *Arabidopsis* is predominantly selfing. For variants with alternate allele frequency greater than 50%, we flipped the sign of GPN scores (equivalent to taking the log-likelihood ratio between the minor and the major allele), and did all analyses in terms of minor allele frequency. Variant consequences produced by Ensembl Variant Effect Predictor were downloaded from Ensembl Plants. Conservation scores were downloaded from http://plantregmap.gao-lab.org/download.php#alignment-conservation. For conservation scores phyloP and phastCons, we simply flipped the sign to obtain a variant score, i.e., variants at conserved sites should be considered more pathogenic. We additionally scored variants using (minus) the absolute value of phyloP, referred to as abs(phyloP), which means prioritizing putative accelerated regions over putative neutral ones. We defined rare variants as those with allele count equal to 1 (to be precise, it is two alleles in the same homozygous accession), and common variants as those with allele frequency above 5%. Model scores were defined as pathogenic or benign based on a quantile threshold that we varied from 0.1% to 10%.

GWAS summary statistics were downloaded through the AraGWAS API, with the default threshold of minimum allele count of 6 (i.e., at least 6 homozygous accessions having the allele). The summary statistics include information on whether an association is significant according to a permutation-based approach (recommended [43]) as well as a Bonferroni threshold. The LD matrix of squared Pearson correlations (*r*^2^) was calculated within a radius of 100 kb around each variant, using sgkit (https://pystatgen.github.io/). We define a weighted sum of GPN scores according to LD (*i* and *j* index SNPs):

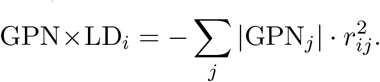

This is an unsigned version of the Signed LD Profile [57] and can also be interpreted as the multiplication between the LD matrix and the vector of GPN scores. The reason why we used unsigned LD and model scores is that we focused on assessing whether a variant would have a significant association with differences in a trait, regardless of the direction of the association. Since the association *p*-value is invariant to recoding of reference and alternate alleles, we took the absolute value of GPN scores. We arbitrarily added a negative sign in front to be consistent with more negative implying more likely functional. We similarly defined phyloP*×*LD (which first required shifting the scores to the negative half-line), abs(phyloP)*×* and phastCons*×*LD. We considered the baseline LD Score [58], the unweighted sum of LD with a given variant:

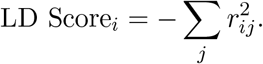

## Code availability

Code to reproduce all results, including instructions to load the pre-trained model, is available at https://github.com/songlab-cal/gpn.

## Use of AI software

ChatGPT was used to improve the wording of some paragraphs, but not to generate new content.

## Acknowledgments

We would like to thank Carlos Albors, Jesús Martínez-Gómez, Eyes Robson, Nilah Ioannidis and Allison Gaudinier for helpful discussions. This research is supported in part by an NIH grant R35-GM134922 and a grant from the Koret-UC Berkeley-Tel Aviv University Initiative in Computational Biology and Bioinformatics.

## Footnotes

1 https://useast.ensembl.org/info/genome/variation/prediction/predicted_data.html

