## Supplementary Tables ad Figures for "DNA language models are powerful predictors of genome-wide variant effects"

### Supporting Information for
DNA language models are powerful predictors of genome-wide variant effects

Gonzalo Benegas[1], Sanjit Singh Batra[2], and Yun S. Song[2,3,*]

[1] Graduate Group in Computational Biology, University of California, Berkeley
[2] Computer Science Division, University of California, Berkeley
[3] Department of Statistics, University of California, Berkeley

This PDF file includes:

- Tables S1-S3
- Figures S1 to S11

#### Supplementary Tables

**Table S1:** Genome assemblies used for training

| Assembly Accession | Assembly Name | Organism Name |
|---|---|---|
| GCF_000001735.4 | TAIR10.1 | *Arabidopsis thaliana* |
| GCF_000309985.2 | CAAS_Brap_v3.01 | *Brassica rapa* |
| GCF_000633955.1 | Cs | *Camelina sativa* |
| GCF_000375325.1 | Caprub1_0 | *Capsella rubella* |
| GCF_000150535.2 | Papaya1.0 | *Carica papaya* |
| GCF_000478725.1 | Eutsalg1_0 | *Eutrema salsugineum* |
| GCF_000801105.1 | Rs1.0 | *Raphanus sativus* |
| GCF_000463585.1 | ASM46358v1 | *Tarenaya hassleriana* |

**Table S2:** Test perplexity. Perplexity, defined as the exponentiation of the cross-entropy loss, is equivalent to 1 over the probability given to the correct nucleotide. *Arabidopsis thaliana* chromosomes 4 and 5 were used for validation and testing, respectively. Note that reducing the repeat weight leads to improved test perplexity in non-repetitive regions, which are often of greater interest. Compared to full down-weighting, moderate down-weighting results in a similar improvement in perplexity for non-repetitive regions without sacrificing genome-wide perplexity as much.

| Model | Chromosome-wide | Non-repeat regions |
|---|---|---|
| Repeat weight 1 | 2.88 | 2.99 |
| Repeat weight 0.1 | 2.90 | 2.92 |
| Repeat weight 0 | 3.03 | 2.92 |

**Table S3:** Training hyperparameters

| | |
|---|---|
| Window size (L) | 512 |
| Repeat weight | 0.1 |
| Embedding dimension (D) | 512 |
| Convolutional blocks | 25 |
| Convolutional kernel size | 9 |
| Convolutional dilation schedule | $1, 2, 4, 8, 16, 32, 1, 2, 4, 8, 16, 32, \ldots$ |
| Optimizer | AdamW |
| Weight decay | 0.01 |
| Batch size | 2048 |
| Learning rate | $10^{-3}$ for 120 K steps + decaying (cosine) for 30 K steps |
| Learning rate warmup | 1 K steps |

#### Supplementary Figures

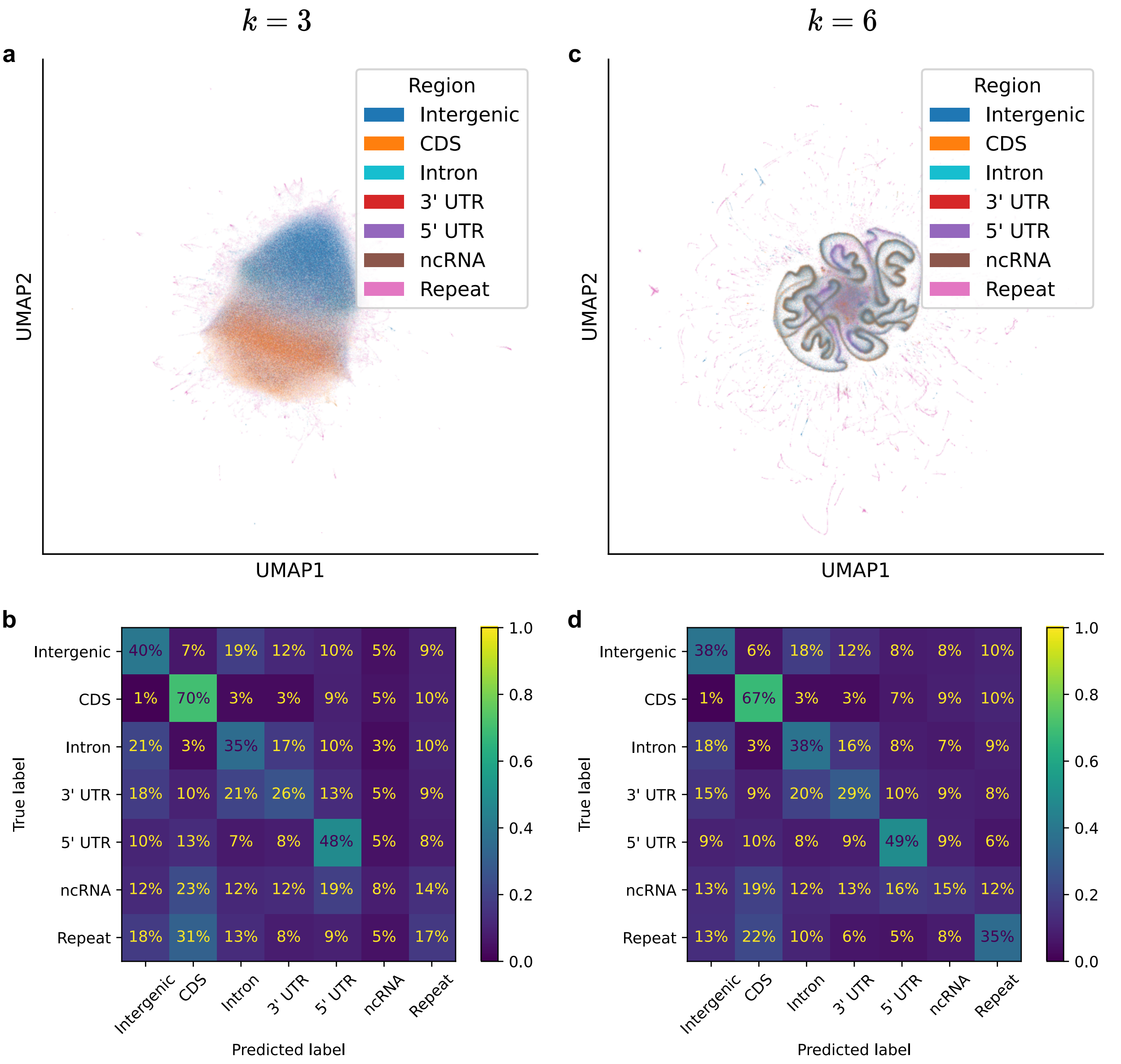


**Figure S1: UMAP visualization of $k$-mer spectrum of different windows, as in Figure 2, annotated with gene region.** (a,b) $k = 3$. (c,d) $k = 6$.

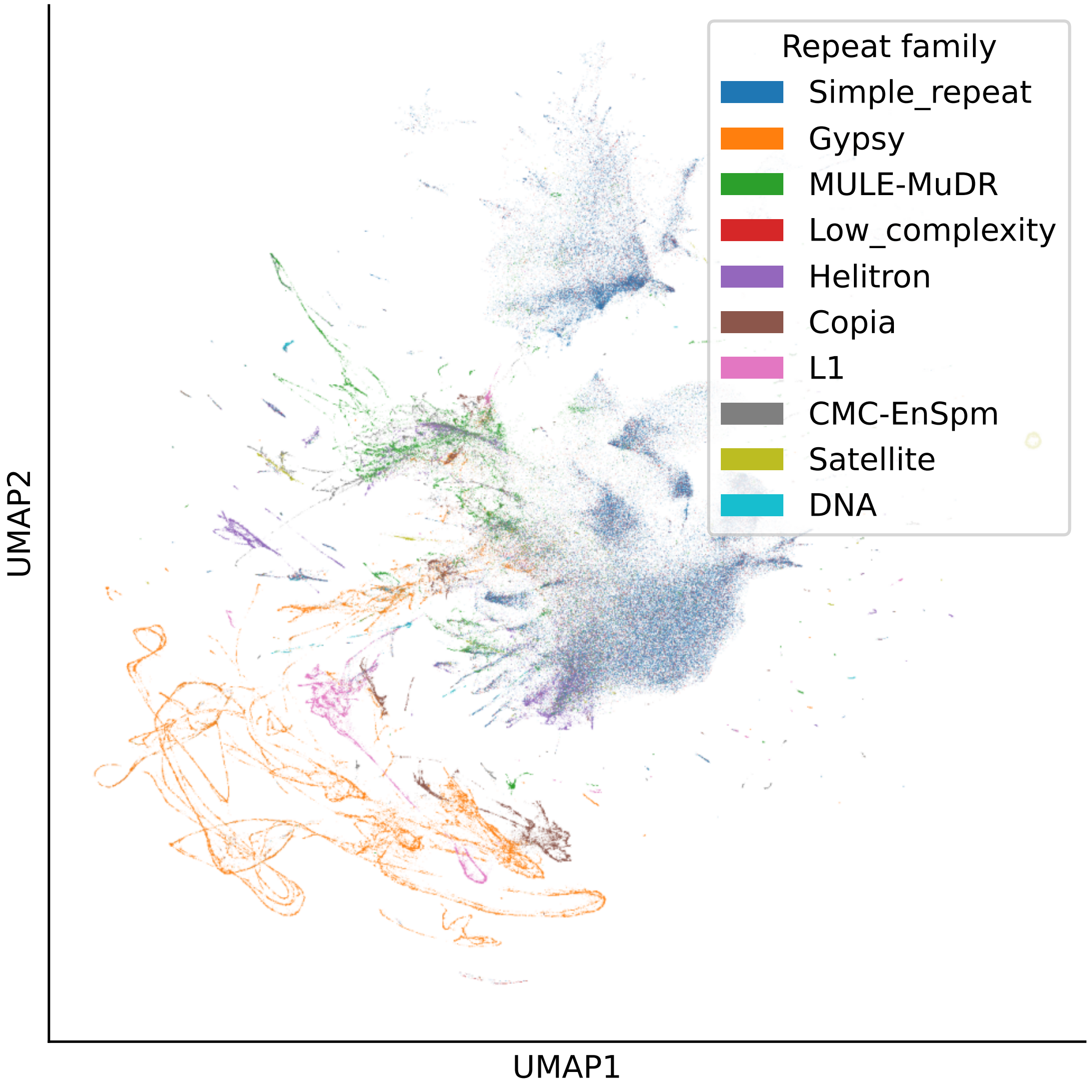


**Figure S2: UMAP visualization of GPN embeddings, as in Figure 2, annotated by repeat family.**

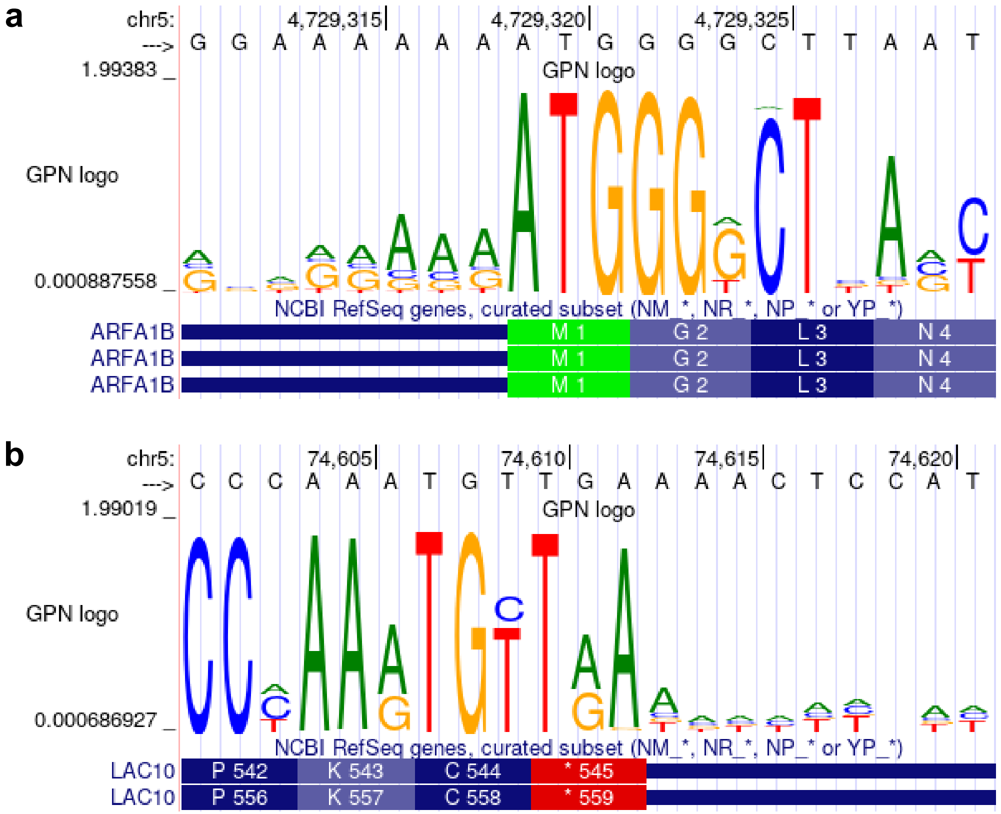


**Figure S3: Additional GPN sequence logos.** (a) Start codon. (b) Stop codon.

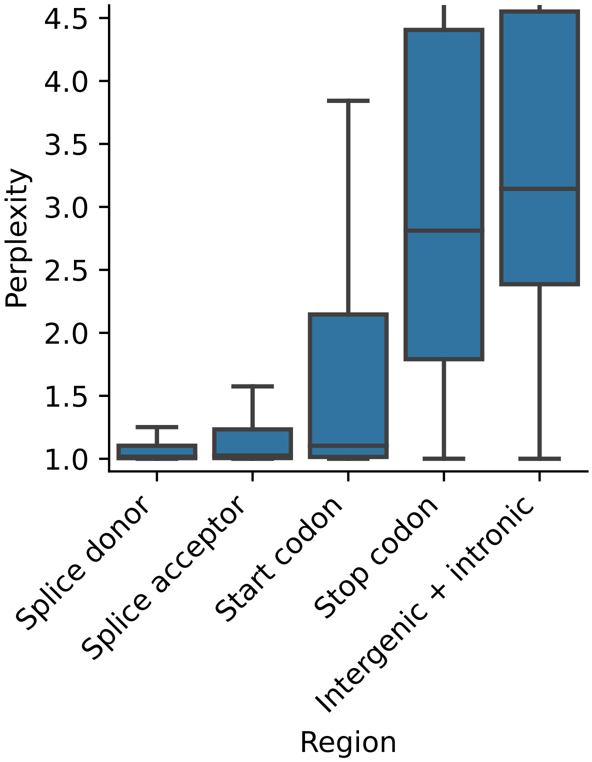


**Figure S4: Perplexity on select regions from the test chromosome.**

| pattern | num_seqlets | modisco_cwm_fwd | modisco_cwm_rev | match0 | qval0 | match0_logo |
|---|---|---|---|---|---|---|
| pos_patterns.pattern_0 | 5028 | | | | | |
| pos_patterns.pattern_1 | 4509 | | | AT4G38000 | 0.0 | |
| pos_patterns.pattern_2 | 3386 | | | AT1G72740 | 0.000117 | |
| pos_patterns.pattern_3 | 1658 | | | | | |
| pos_patterns.pattern_4 | 1611 | | | | | |
| pos_patterns.pattern_5 | 1556 | | | | | |
| pos_patterns.pattern_6 | 1490 | | | AT2G01930 | 0.0 | |
| pos_patterns.pattern_7 | 1424 | | | | | |
| pos_patterns.pattern_8 | 1391 | | | AT5G18090 | 0.044918 | |
| pos_patterns.pattern_9 | 1385 | | | | | |
| pos_patterns.pattern_10 | 1057 | | | | | |
| pos_patterns.pattern_11 | 1052 | | | | | |
| pos_patterns.pattern_12 | 928 | | | | | |
| pos_patterns.pattern_13 | 921 | | | | | |
| pos_patterns.pattern_14 | 844 | | | | | |
| pos_patterns.pattern_15 | 837 | | | | | |
| pos_patterns.pattern_16 | 836 | | | AT3G48430 | 0.000584 | |
| pos_patterns.pattern_17 | 828 | | | | | |

**Figure S5: Promoter motifs predicted by GPN and matching motifs in PlantTFDB.**

| pattern | num_seqlets | modisco_cwm_fwd | modisco_cwm_rev | match0 | qval0 | match0_logo |
|---|---|---|---|---|---|---|
| pos_patterns.pattern_18 | 780 | 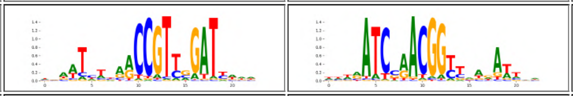 | | AT4G24470 | 0.0 | 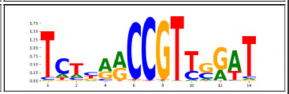 |
| pos_patterns.pattern_19 | 775 | 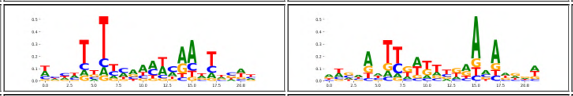 | | AT2G41835 | 0.016114 | 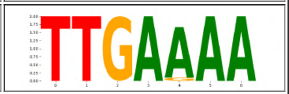 |
| pos_patterns.pattern_20 | 756 | 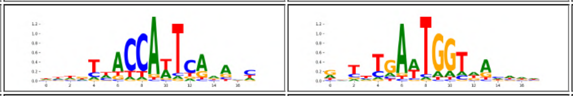 | | | | |
| pos_patterns.pattern_21 | 743 | 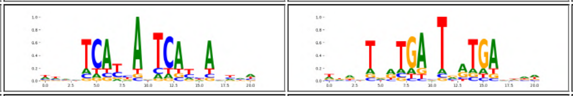 | | | | |
| pos_patterns.pattern_22 | 729 | 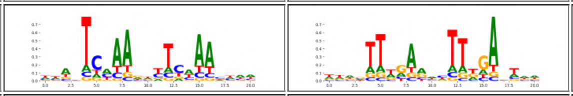 | | AT1G49560 | 0.029145 | 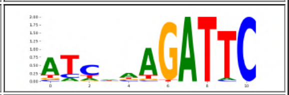 |
| pos_patterns.pattern_23 | 708 | 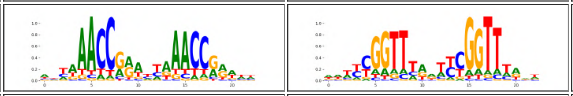 | | | | |
| pos_patterns.pattern_24 | 702 | 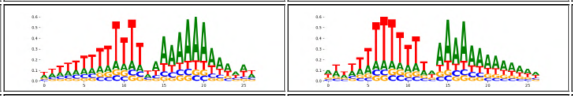 | | | | |
| pos_patterns.pattern_25 | 696 | 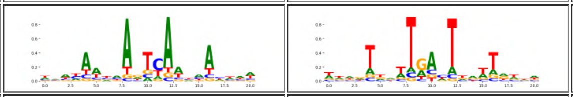 | | | | |
| pos_patterns.pattern_26 | 688 | 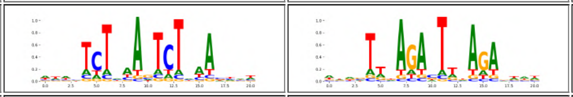 | | | | |
| pos_patterns.pattern_27 | 674 | 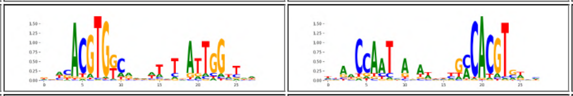 | | | | |
| pos_patterns.pattern_28 | 643 | 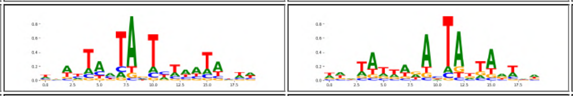 | | AT2G36610 | 0.026016 | 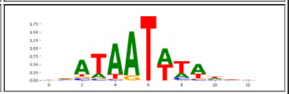 |
| pos_patterns.pattern_29 | 640 | 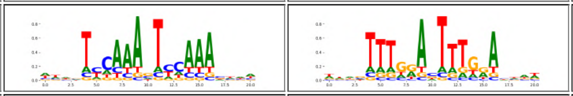 | | | | |
| pos_patterns.pattern_30 | 631 | 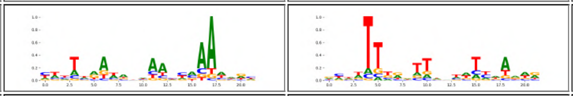 | | | | |
| pos_patterns.pattern_31 | 631 | 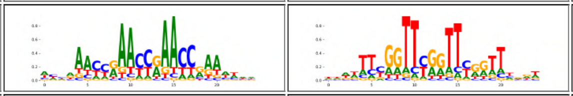 | | | | |
| pos_patterns.pattern_32 | 630 | 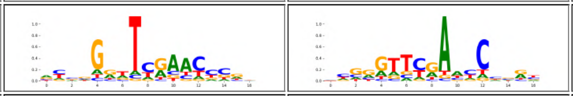 | | | | |
| pos_patterns.pattern_33 | 604 | 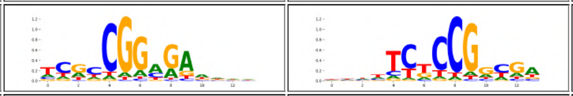 | | AT3G58630 | 0.029229 | 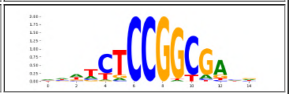 |
| pos_patterns.pattern_34 | 595 | 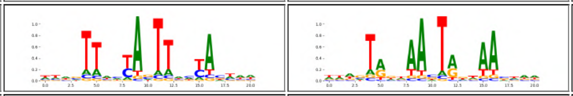 | | | | |
| pos_patterns.pattern_35 | 595 | 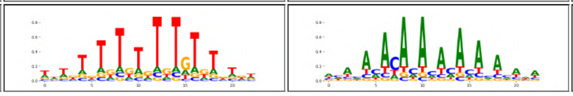 | | | | |

**Figure S4:** (Continued)

| pattern | num_seqlets | modisco_cwm_fwd | modisco_cwm_rev | match0 | qval0 | match0_logo |
|---|---|---|---|---|---|---|
| pos_patterns.pattern_36 | 590 | 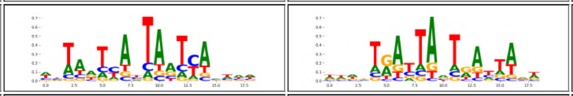 | | AT3G28920 | 0.000035 | 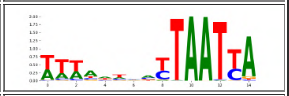 |
| pos_patterns.pattern_37 | 580 | 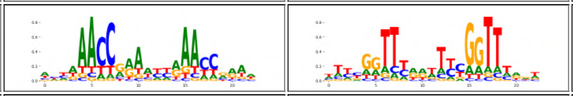 | | | | |
| pos_patterns.pattern_38 | 566 | 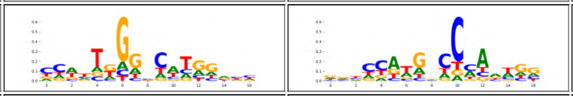 | | | | |
| pos_patterns.pattern_39 | 558 | 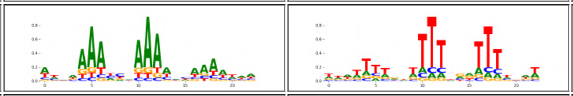 | | | | |
| pos_patterns.pattern_40 | 544 | 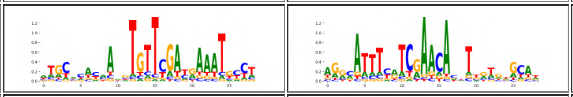 | | | | |
| pos_patterns.pattern_41 | 523 | 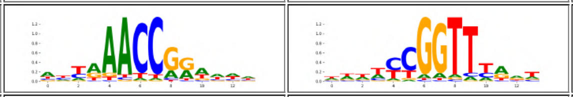 | | | | |
| pos_patterns.pattern_42 | 512 | 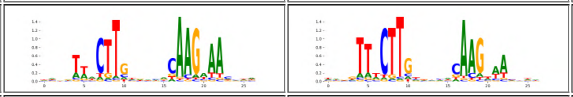 | | AT3G10480 | 0.000001 | 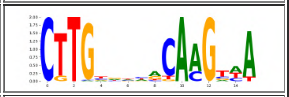 |
| pos_patterns.pattern_43 | 511 | 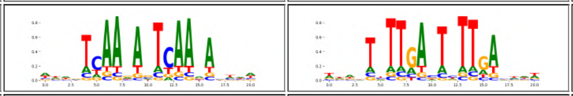 | | | | |
| pos_patterns.pattern_44 | 508 | 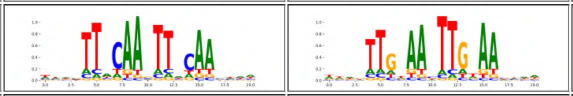 | | | | |
| pos_patterns.pattern_45 | 506 | 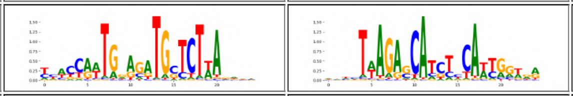 | | | | |
| pos_patterns.pattern_46 | 492 | 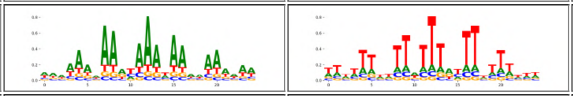 | | AT2G28810 | 0.000187 | 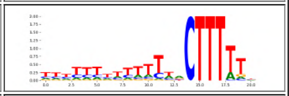 |
| pos_patterns.pattern_47 | 480 | 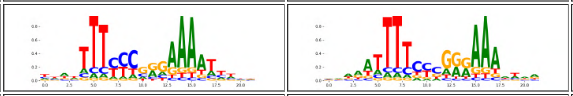 | | | | |
| pos_patterns.pattern_48 | 480 | 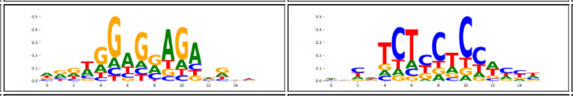 | | | | |
| pos_patterns.pattern_49 | 477 | 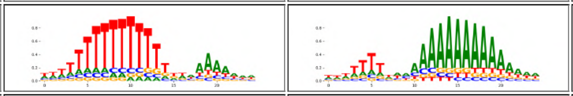 | | AT4G38000 | 0.002972 | 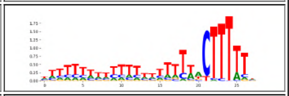 |
| pos_patterns.pattern_50 | 454 | 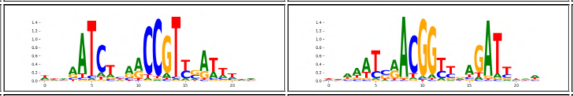 | | AT4G24470 | 0.000033 | 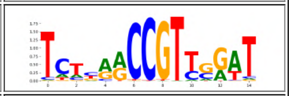 |
| pos_patterns.pattern_51 | 451 | 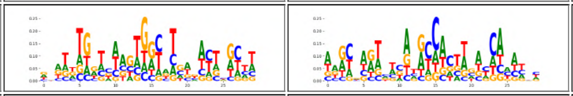 | | AT3G10800 | 0.025896 | 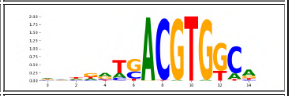 |
| pos_patterns.pattern_52 | 433 | 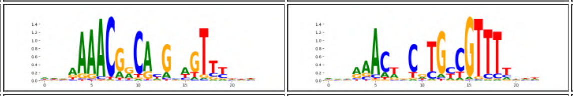 | | | | |
| pos_patterns.pattern_53 | 426 | 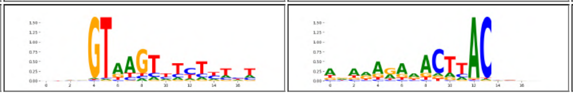 | | | | |

**Figure S4:** (Continued)

| pattern | num_seqlets | modisco_cwm_fwd | modisco_cwm_rev | match0 | qval0 | match0_logo |
|---|---|---|---|---|---|---|
| pos_patterns.pattern_54 | 407 | | | AT2G01930 | 0.002768 | |
| pos_patterns.pattern_55 | 404 | | | | | |
| pos_patterns.pattern_56 | 387 | | | AT4G34000 | 0.000455 | |
| pos_patterns.pattern_57 | 357 | | | AT5G42520 | 0.037446 | |
| pos_patterns.pattern_58 | 354 | | | AT1G69570 | 0.0 | |
| pos_patterns.pattern_59 | 352 | | | AT3G62420 | 0.000053 | |
| pos_patterns.pattern_60 | 322 | | | | | |
| pos_patterns.pattern_61 | 319 | | | | | |
| pos_patterns.pattern_62 | 314 | | | | | |
| pos_patterns.pattern_63 | 311 | | | | | |
| pos_patterns.pattern_64 | 309 | | | | | |
| pos_patterns.pattern_65 | 302 | | | AT1G21910 | 0.0 | |
| pos_patterns.pattern_66 | 293 | | | | | |
| pos_patterns.pattern_67 | 291 | | | AT2G33860 | 0.005094 | |
| pos_patterns.pattern_68 | 285 | | | | | |
| pos_patterns.pattern_69 | 282 | | | | | |
| pos_patterns.pattern_70 | 280 | | | | | |
| pos_patterns.pattern_71 | 280 | | | AT1G53170 | 0.001163 | |

**Figure S4:** (Continued)

| pattern | num_seqlets | modisco_cwm_fwd | modisco_cwm_rev | match0 | qval0 | match0_logo |
|---|---|---|---|---|---|---|
| pos_patterns.pattern_72 | 275 | 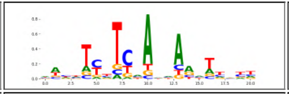 | 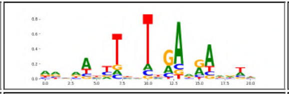 | | | |
| pos_patterns.pattern_73 | 265 | 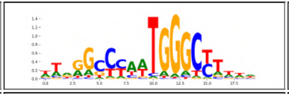 | 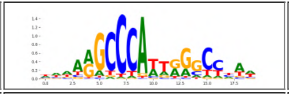 | | | |
| pos_patterns.pattern_74 | 263 | 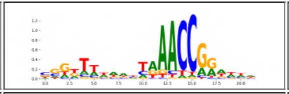 | 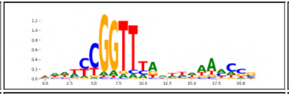 | | | |
| pos_patterns.pattern_75 | 262 | 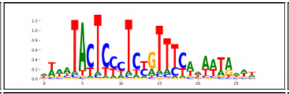 | 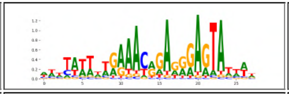 | | | |
| pos_patterns.pattern_76 | 259 | 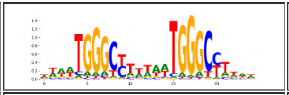 | 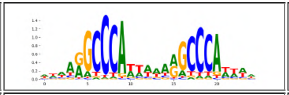 | | | |
| pos_patterns.pattern_77 | 253 | 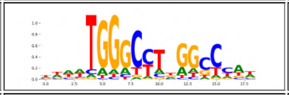 | 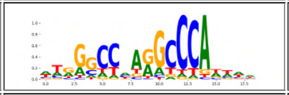 | | | |
| pos_patterns.pattern_78 | 249 | 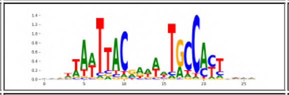 | 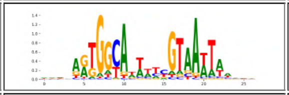 | | | |
| pos_patterns.pattern_79 | 248 | 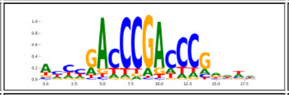 | 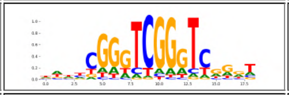 | | | |
| pos_patterns.pattern_80 | 242 | 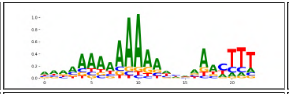 | 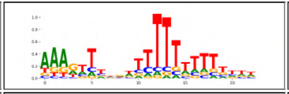 | AT1G49480 | 0.000331 | 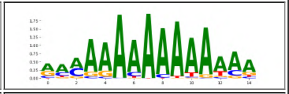 |
| pos_patterns.pattern_81 | 228 | 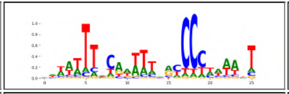 | 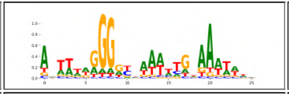 | AT5G67580 | 0.016528 | 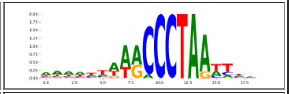 |
| pos_patterns.pattern_82 | 225 | 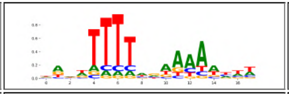 | 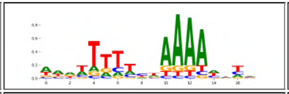 | | | |
| pos_patterns.pattern_83 | 221 | 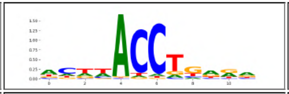 | 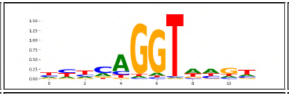 | | | |
| pos_patterns.pattern_84 | 215 | 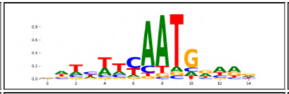 | 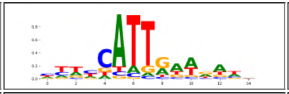 | | | |
| pos_patterns.pattern_85 | 209 | 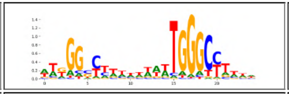 | 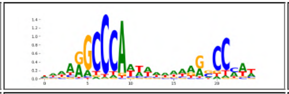 | | | |
| pos_patterns.pattern_86 | 206 | 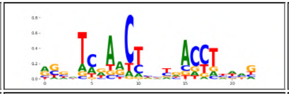 | 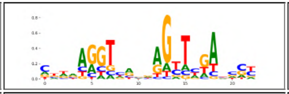 | | | |
| pos_patterns.pattern_87 | 199 | 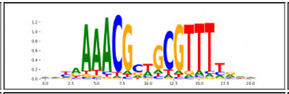 | 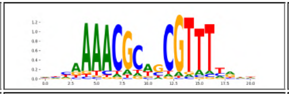 | | | |
| pos_patterns.pattern_88 | 198 | 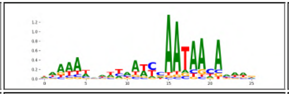 | 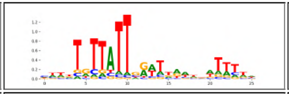 | | | |
| pos_patterns.pattern_89 | 194 | 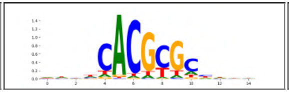 | 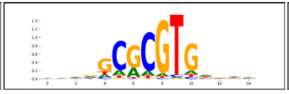 | AT3G22170 | 0.002435 | 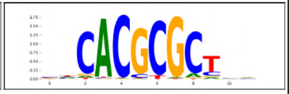 |

**Figure S4:** (Continued)

| pattern | num_seqlets | modisco_cwm_fwd | modisco_cwm_rev | match0 | qval0 | match0_logo |
|---|---|---|---|---|---|---|
| pos_patterns.pattern_90 | 186 | 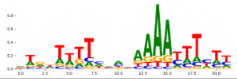 | 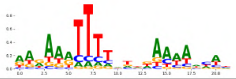 | | | |
| pos_patterns.pattern_91 | 169 | 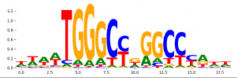 | 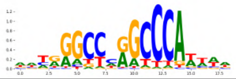 | | | |
| pos_patterns.pattern_92 | 167 | 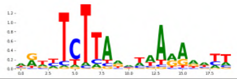 | 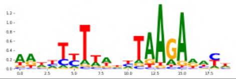 | | | |
| pos_patterns.pattern_93 | 167 | 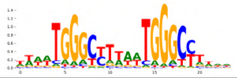 | 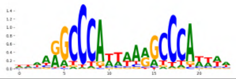 | | | |
| pos_patterns.pattern_94 | 167 | 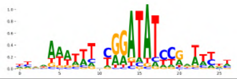 | 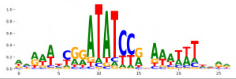 | | | |
| pos_patterns.pattern_95 | 159 | 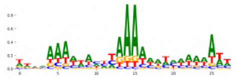 | 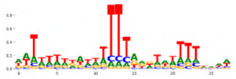 | AT4G38000 | 0.005238 | 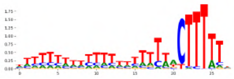 |
| pos_patterns.pattern_96 | 158 | 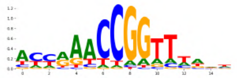 | 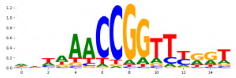 | | | |
| pos_patterns.pattern_97 | 157 | 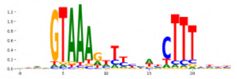 | 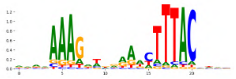 | | | |
| pos_patterns.pattern_98 | 157 | 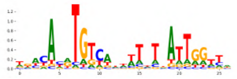 | 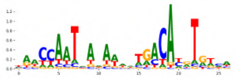 | | | |
| pos_patterns.pattern_99 | 156 | 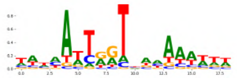 | 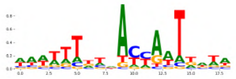 | | | |
| pos_patterns.pattern_100 | 154 | 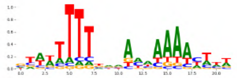 | 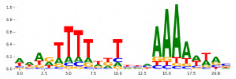 | | | |
| pos_patterns.pattern_101 | 151 | 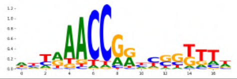 | 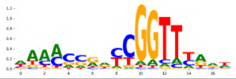 | | | |
| pos_patterns.pattern_102 | 146 | 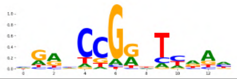 | 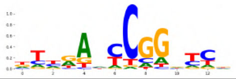 | | | |
| pos_patterns.pattern_103 | 136 | 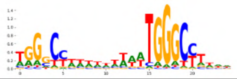 | 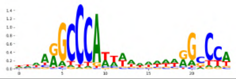 | | | |
| pos_patterns.pattern_104 | 135 | 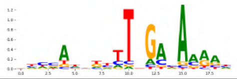 | 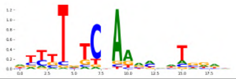 | | | |
| pos_patterns.pattern_105 | 135 | 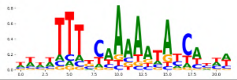 | 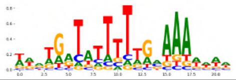 | AT5G02840 | 0.041659 | 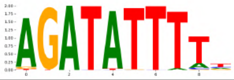 |
| pos_patterns.pattern_106 | 131 | 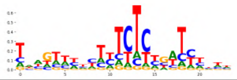 | 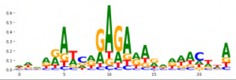 | | | |
| pos_patterns.pattern_107 | 131 | 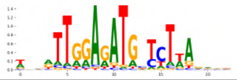 | 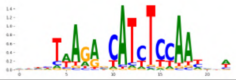 | | | |

**Figure S4:** (Continued)

| pattern | num_seqlets | modisco_cwm_fwd | modisco_cwm_rev | match0 | qval0 | match0_logo |
|---|---|---|---|---|---|---|
| pos_patterns.pattern_108 | 130 | | | | | |
| pos_patterns.pattern_109 | 128 | | | | | |
| pos_patterns.pattern_110 | 126 | | | | | |
| pos_patterns.pattern_111 | 125 | | | | | |
| pos_patterns.pattern_112 | 121 | | | | | |
| pos_patterns.pattern_113 | 117 | | | | | |
| pos_patterns.pattern_114 | 115 | | | | | |
| pos_patterns.pattern_115 | 107 | | | AT2G20110 | 0.049803 | |
| pos_patterns.pattern_116 | 106 | | | | | |
| pos_patterns.pattern_117 | 105 | | | | | |
| pos_patterns.pattern_118 | 105 | | | AT3G55370 | 0.025597 | |
| pos_patterns.pattern_119 | 104 | | | | | |
| pos_patterns.pattern_120 | 103 | | | | | |
| pos_patterns.pattern_121 | 102 | | | | | |
| pos_patterns.pattern_122 | 100 | | | AT3G22170 | 0.007859 | |
| pos_patterns.pattern_123 | 96 | | | | | |
| pos_patterns.pattern_124 | 93 | | | AT4G24470 | 0.000435 | |
| pos_patterns.pattern_125 | 91 | | | AT3G10500 | 0.000013 | |

**Figure S4:** (Continued)

| pattern | num_seqlets | modisco_cwm_fwd | modisco_cwm_rev | match0 | qval0 | match0_logo |
|---|---|---|---|---|---|---|
| pos_patterns.pattern_126 | 88 | 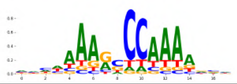 | 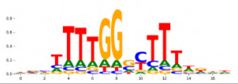 | | | |
| pos_patterns.pattern_127 | 88 | 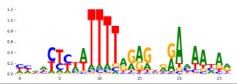 | 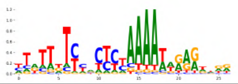 | | | |
| pos_patterns.pattern_128 | 87 | 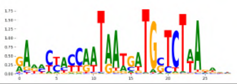 | 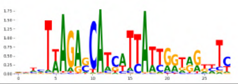 | | | |
| pos_patterns.pattern_129 | 86 | 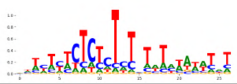 | 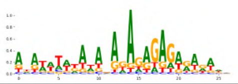 | AT2G01930 | 0.007347 | 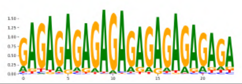 |
| pos_patterns.pattern_130 | 80 | 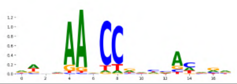 | 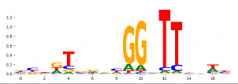 | | | |
| pos_patterns.pattern_131 | 79 | 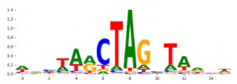 | 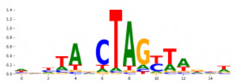 | | | |
| pos_patterns.pattern_132 | 75 | 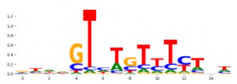 | 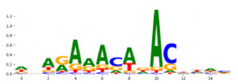 | | | |
| pos_patterns.pattern_133 | 72 | 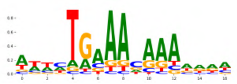 | 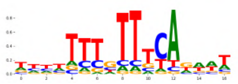 | | | |
| pos_patterns.pattern_134 | 67 | 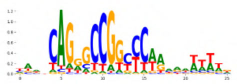 | 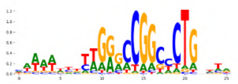 | | | |
| pos_patterns.pattern_135 | 63 | 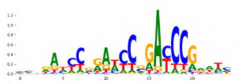 | 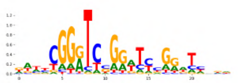 | | | |
| pos_patterns.pattern_136 | 61 | 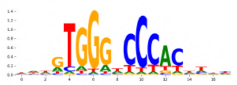 | 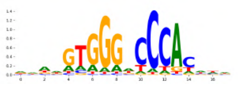 | AT5G23280 | 0.000262 | 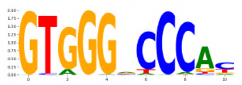 |
| pos_patterns.pattern_137 | 61 | 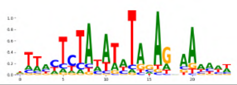 | 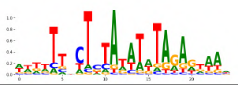 | | | |
| pos_patterns.pattern_138 | 61 | 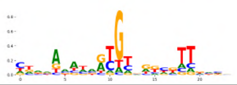 | 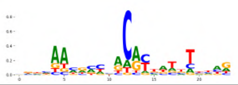 | | | |
| pos_patterns.pattern_139 | 56 | 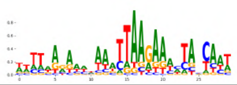 | 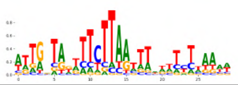 | | | |
| pos_patterns.pattern_140 | 55 | 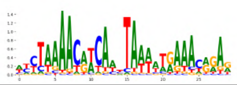 | 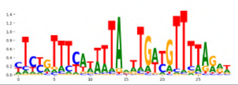 | | | |
| pos_patterns.pattern_141 | 54 | 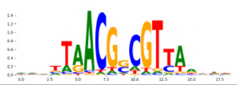 | 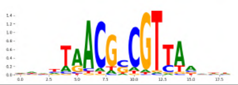 | AT3G10030 | 0.000039 | 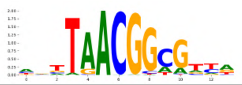 |
| pos_patterns.pattern_142 | 52 | 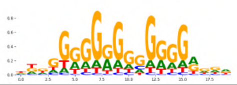 | 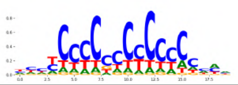 | AT1G67260 | 0.031614 | 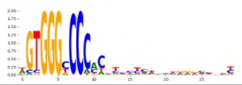 |
| pos_patterns.pattern_143 | 52 | 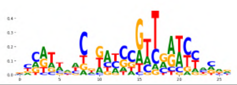 | 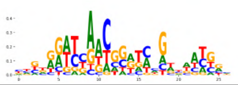 | | | |

**Figure S4:** (Continued)

| pattern | num_seqlets | modisco_cwm_fwd | modisco_cwm_rev | match0 | qval0 | match0_logo |
|---|---|---|---|---|---|---|
| pos_patterns.pattern_144 | 49 | 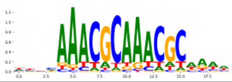 | 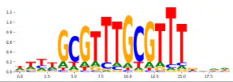 | | | |
| pos_patterns.pattern_145 | 49 | 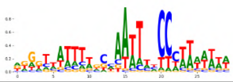 | 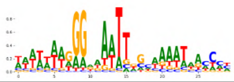 | | | |
| pos_patterns.pattern_146 | 45 | 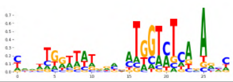 | 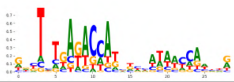 | | | |
| pos_patterns.pattern_147 | 41 | 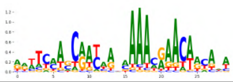 | 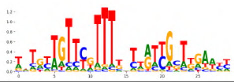 | | | |
| pos_patterns.pattern_148 | 40 | 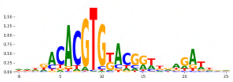 | 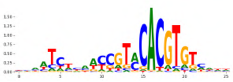 | AT4G34000 | 0.002944 | 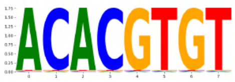 |
| pos_patterns.pattern_149 | 39 | 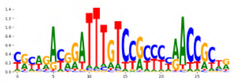 | 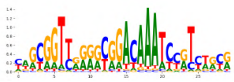 | | | |
| pos_patterns.pattern_150 | 35 | 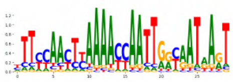 | 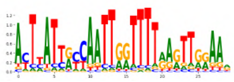 | | | |
| pos_patterns.pattern_151 | 35 | 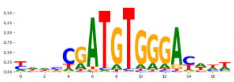 | 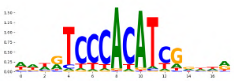 | | | |
| pos_patterns.pattern_152 | 32 | 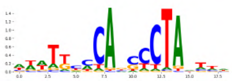 | 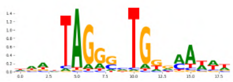 | | | |
| pos_patterns.pattern_153 | 29 | 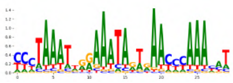 | 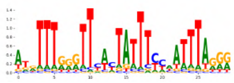 | | | |
| pos_patterns.pattern_154 | 28 | 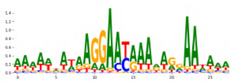 | 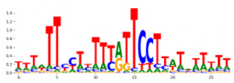 | AT2G45660 | 0.02262 | 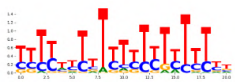 |
| pos_patterns.pattern_155 | 28 | 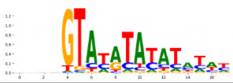 | 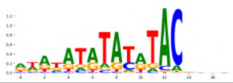 | | | |
| pos_patterns.pattern_156 | 28 | 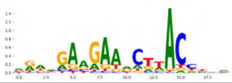 | 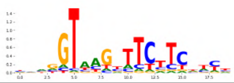 | | | |
| pos_patterns.pattern_157 | 27 | 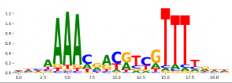 | 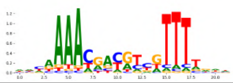 | | | |
| pos_patterns.pattern_158 | 27 | 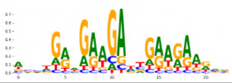 | 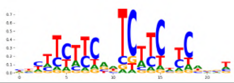 | | | |
| pos_patterns.pattern_159 | 26 | 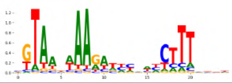 | 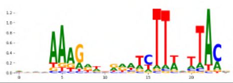 | | | |
| pos_patterns.pattern_160 | 26 | 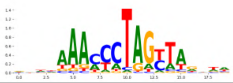 | 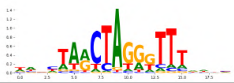 | | | |
| pos_patterns.pattern_161 | 23 | 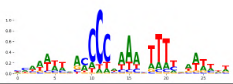 | 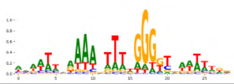 | | | |

**Figure S4:** (Continued)

| pattern | num_seqlets | modisco_cwm_fwd | modisco_cwm_rev | match0 | qval0 | match0_logo |
|---|---|---|---|---|---|---|
| pos_patterns.pattern_162 | 23 | 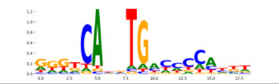 | 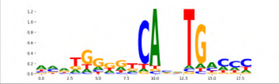 | | | |
| pos_patterns.pattern_163 | 22 | 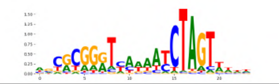 | 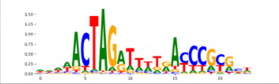 | | | |

**Figure S4:** (Continued)

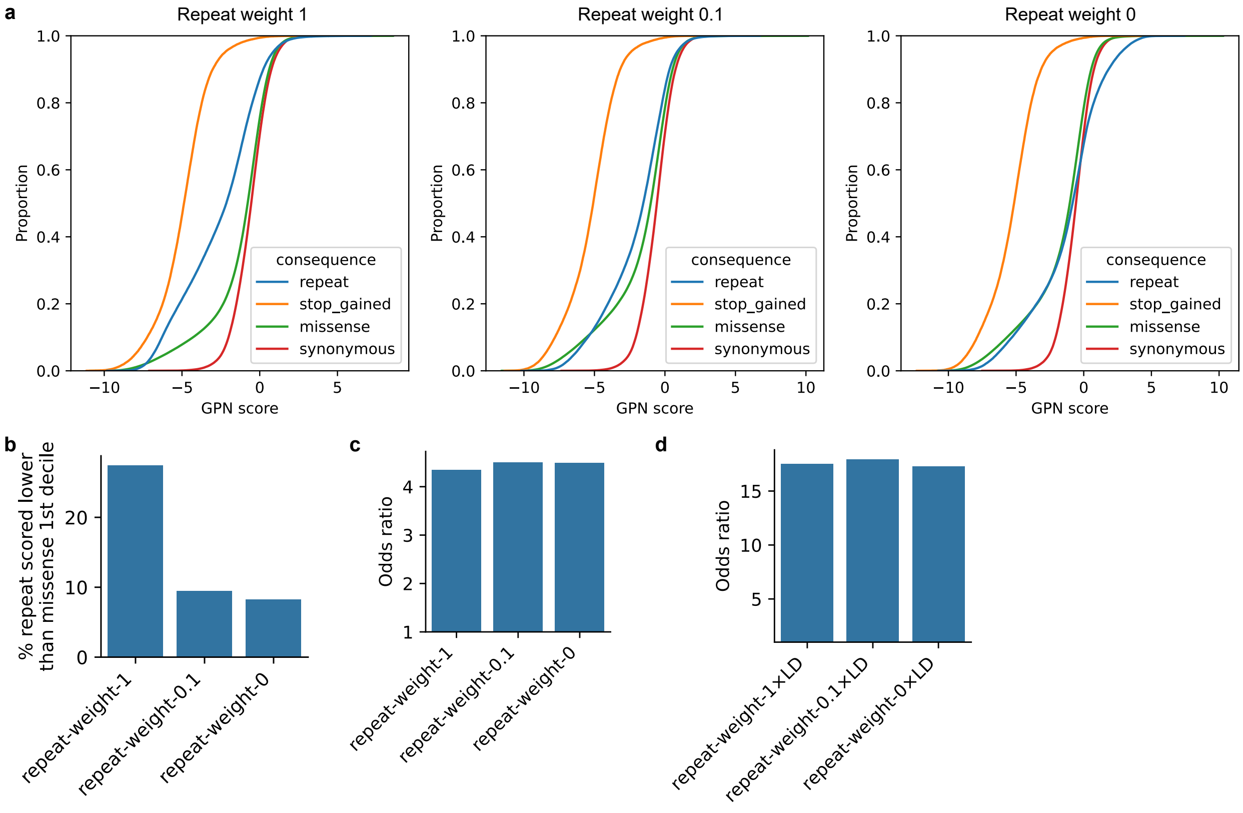


**Figure S5: Comparison of models trained with different loss weights on repeats.** (a) Cumulative distribution function of GPN scores for simulated variants in specific categories, as described in Figure 4. (b) Percentage of simulated repeat variants scored lower than the first decile of simulated missense variants. (c) Odds ratios for rare vs. common variants, as described in Figure 5c. (d) Odds ratios for GWAS hits, as described in Figure 6c.

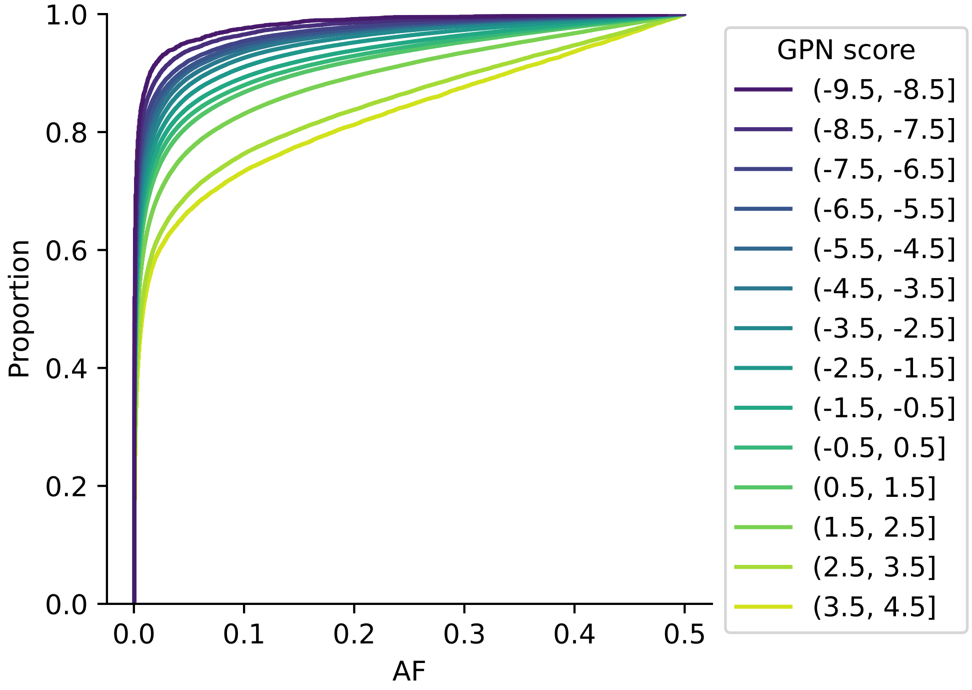


**Figure S6: Cumulative distribution function of allele frequency (AF) for variants in different GPN score bins, as described in Figure 5b.**

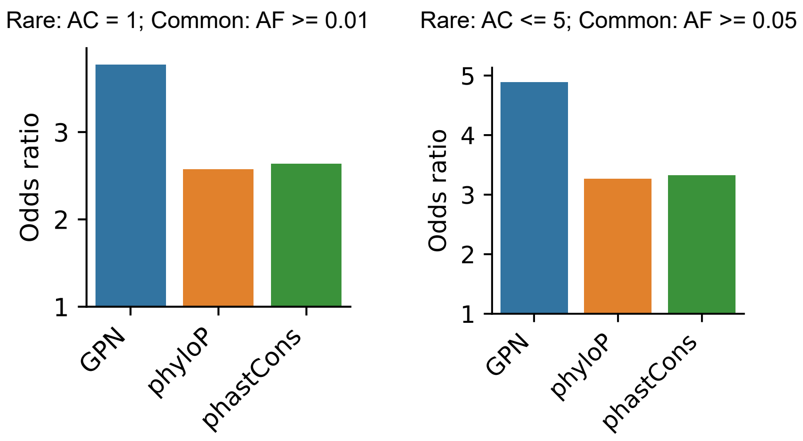


**Figure S7: Rare vs. common odds ratios for different thresholds for defining rare and common variants.** Odds ratios (OR) were calculated as described in Figure 5c.

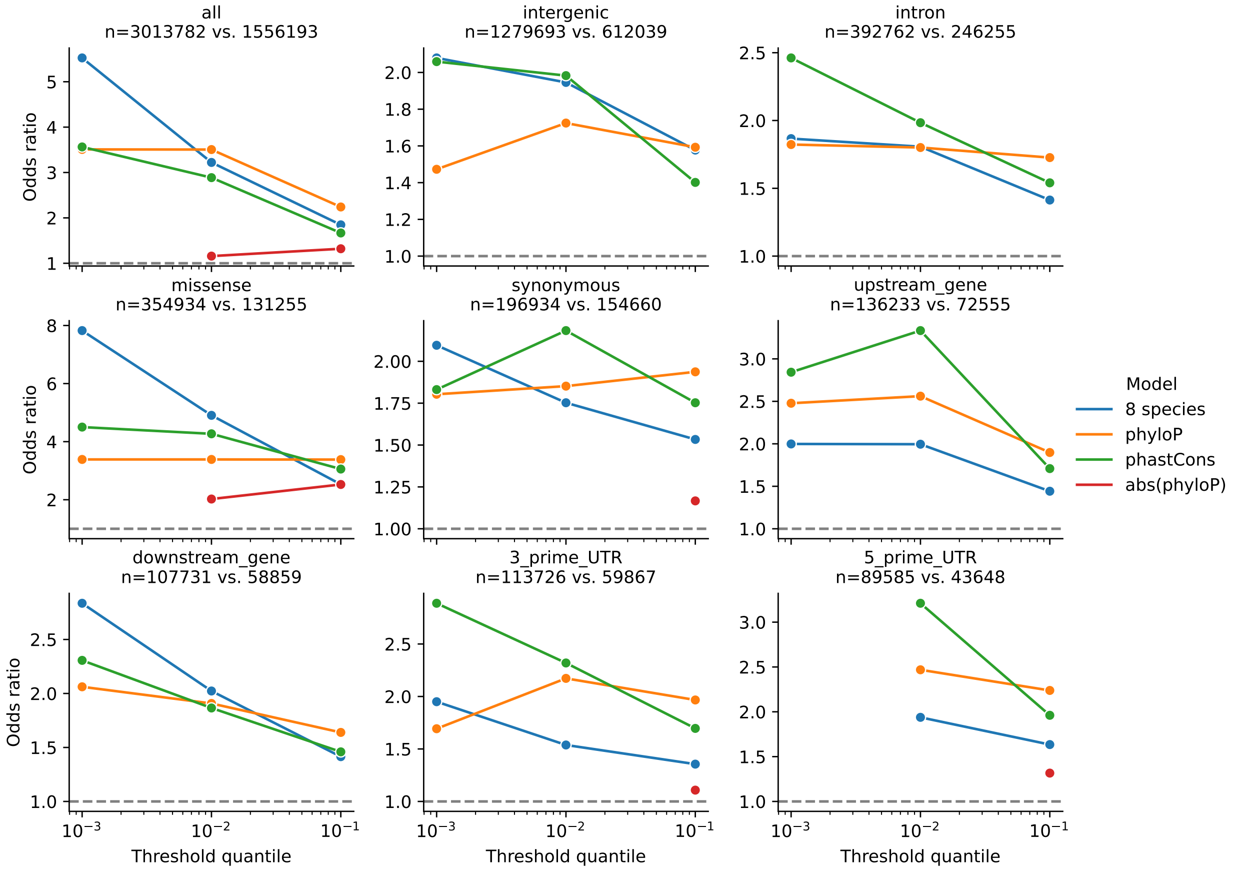


**Figure S8: Rare vs. common odds ratios for specific variant categories and different thresholds for defining functional scores.** Odds ratios (OR) were calculated as described in Figure 5c. Only significant odds ratios are shown. The most stringent threshold in 5' UTR was excluded due to certain models having less than 10 counts in an entry of the contingency table.

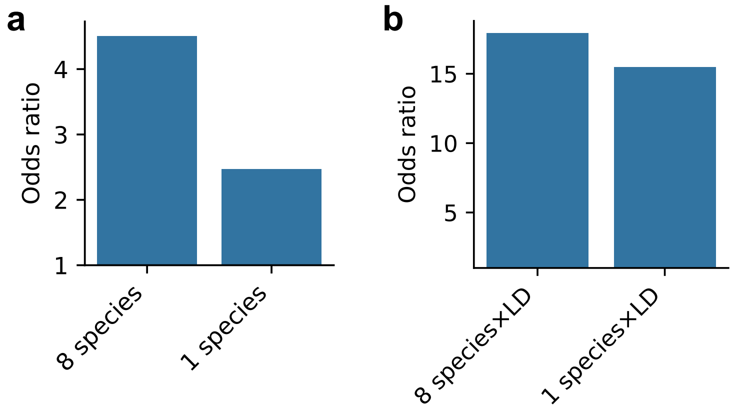


**Figure S9: Comparison of models trained on a different number of species.** (a) Odds ratios for rare vs. common variants, as described in Figure 5c. (b) Odds ratios for GWAS hits, as described in Figure 6c.

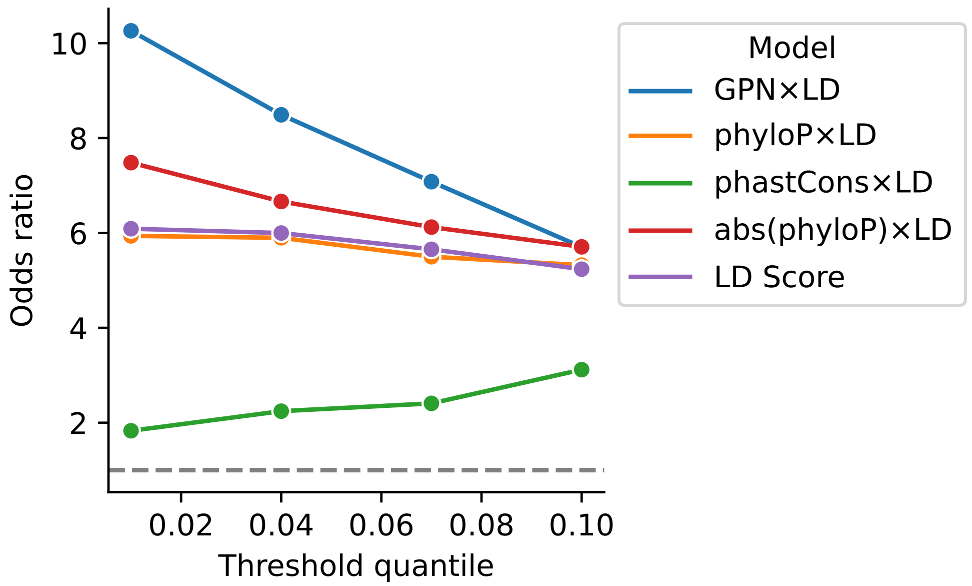


**Figure S10: GWAS hit odds ratios for different thresholds for defining functional-tagged scores.** Odds ratios (OR) were calculated as described in Figure 6c.

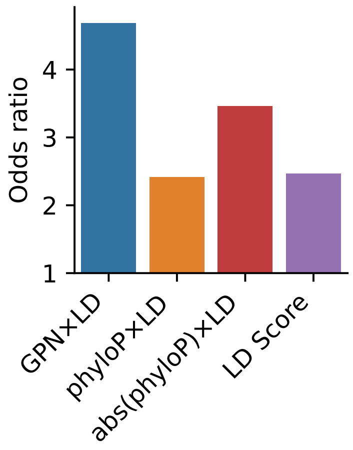


**Figure S11: Odds ratios for GWAS hits, using the Bonferroni correction instead of permutation-based significance threshold, as described in Figure 6c.** Only significant odds ratios are shown.
